# From Intrinsic Topography to Cognitive Prediction: Benchmarking Approaches to Personalized Cortical Parcellation

**DOI:** 10.64898/2026.10.09.758001

**Authors:** Yusuf Cosku Inceler, Micha Burkhardt, Carsten Gießing, Daniel Kristanto

## Abstract

Personalized functional magnetic resonance imaging (fMRI) adapts cortical parcellations to individual functional anatomy. Various methods have been proposed that differ in how they combine group-level information, spatial constraints, and individual functional data. Using the Human Connectome Project Young Adult dataset (*N* = 947), we compared five personalized parcellation methods and a standard group-level atlas.

For each method, we calculated functional homogeneity, test-retest reliability, parcel contiguity, and inter-individual Dice overlap. Personalized parcellation methods generally yielded higher functional homogeneity and lower inter-individual Dice than the group atlas, while test-retest reliability of functional connectivity was high across methods. Parcel contiguity differed across methods: some approaches produced almost entirely contiguous parcels, whereas others frequently produced spatially disconnected components, indicating that the detection of candidate ectopic variants depends partly on the parcellation strategy.

We also examined whether connectivity and topographic features were associated with and predicted general intelligence (*g*). Using structural equation modeling, intra-individual parcel size variability and inter-individual parcel size similarity derived from the hierarchical Bayesian method (gMSHBM) were positively associated with *g*, whereas no corresponding association survived correction for the other methods. Using kernel ridge regression, functional connectivity significantly predicted *g*, with the group atlas yielding the highest predictive accuracy, consistent with a potential advantage of cross-individual spatial correspondence for edge-wise prediction.

Finally, we assessed robustness across four preprocessing pipelines. Although gMSHBM parcellations were not re-estimated, relative between-method patterns in several spatial characteristics were largely preserved. In contrast, brain-behavior associations involving time-series-derived features varied substantially with preprocessing, whereas parcel-size associations were more stable. We conclude that personalized parcellation approaches are associated with distinct trade-offs in spatial correspondence, topographic variability, and downstream predictive utility. The choice of parcellation strategy therefore affects both the properties of individualized maps and the brain-behavior information derived from them.

## 1. Introduction

A central objective of neuroimaging is to map robust brain-behavior associations that explain individual cognitive differences and serve as clinical biomarkers (Finn et al., 2015; Marek et al., 2022). To quantify these relationships, functional Magnetic Resonance Imaging (fMRI) traditionally relies on group-level cortical atlases (Eickhoff et al., 2018). These atlases provide a common coordinate system by assuming that functional organization is spatially consistent across individuals (e.g., Schaefer et al., 2018; Shen et al., 2013; Tzourio-Mazoyer et al., 2002). Precision fMRI studies using densely sampled data challenge this assumption, demonstrating that the spatial layout of functional networks varies substantially across people (Gordon et al., 2017; Laumann et al., 2015). This spatial variability is not uniform; it peaks in association cortices and has been associated with cognitive performance (Cui et al., 2020; Kong et al., 2019). Consequently, personalized parcellation methods have emerged to adapt cortical boundaries to individual functional anatomy (Li et al., 2025).

Personalized functional network mapping reveals two distinct forms of functional variation, namely border shifts and ectopic variants (Dworetsky et al., 2024). Border shifts represent the local displacement of boundaries between neighboring regions. Ectopic variants manifest as spatially disconnected islands separated from the main body of a functional territory. While these phenomena were mapped extensively at the large-scale network level, comparable topological variations also occur at the regional scale. For example, area 55b can split into disconnected components or swap positions with adjacent fields, and area hOc5 can appear in an entirely different sulcus away from its group-average location (Fischl et al., 2008; Yang et al., 2025). These variants potentially reflect neurobiological functional reorganization (Dworetsky et al., 2024), developmental differences (Cui et al., 2020), or compensatory mechanisms (Knights et al., 2025). Moreover, capturing this spatial precision may optimize personalized clinical interventions, such as person-specific therapeutic targeting (Lynch et al., 2022). However, distinguishing true idiosyncratic topography from individual-level noise remains difficult. A highly individualized boundary might capture genuine neurobiological functional displacement, or it might simply overfit motion artifacts and transient signal fluctuations.

This challenge introduces a core methodological problem. Parcellation algorithms differ fundamentally in how they incorporate group-level information, spatial constraints, and individual functional data (Li et al., 2025). Methods applying strict spatial or probabilistic group constraints can preserve cross-individual correspondence but may restrict individual variation. However, they risk masking genuine topological variants. Conversely, methods with fewer or no explicit group-derived constraints can accommodate greater individual spatial variability but may show lower cross-individual correspondence and greater sensitivity to noise. Importantly, these approaches differ in multiple algorithmic properties and should not be interpreted as lying on a single continuum of group constraint or individualization (Li et al., 2025). How these different methodological strategies influence the resulting brain maps and their predictive utility remains uncharacterized. Furthermore, because preprocessing steps such as temporal filtering and global signal regression reshape the functional covariance structures, algorithmic performance cannot be fully characterized under a single analytical pipeline (Botvinik-Nezer et al., 2020; Burkhardt & Gießing, 2026; Kristanto et al., 2024; Luppi et al., 2024; Short et al., 2026; Simmons et al., 2011).

To systematically address this methodological variation, we compared five personalized parcellation methods representing different strategies for incorporating group-level and individual information, against a standard group-level atlas (Schaefer et al., 2018). We leveraged large-scale data from 947 participants in the Human Connectome Project. By evaluating these methods under four distinct preprocessing pipelines, we addressed three specific research questions. First, we evaluated how these different parcellation approaches shape intrinsic map properties, specifically functional homogeneity, test-retest reliability, and spatial contiguity. Second, we investigated how the evaluated parcellation approaches and mathematical distance thresholds alter the detection of ectopic variants. Finally, we assessed how differences in spatial correspondence and personalized topography across methods relate to the out-of-sample prediction of general intelligence (*g*). For the brain-behavior analyses, we focused on widely used features capturing complementary aspects of functional organization: functional connectivity, global network efficiency, and parcel-size topography, which have previously been linked to individual differences in cognition (Kong et al., 2019, 2021; Luppi et al., 2024).

## 2. Methods

### 2.1. Dataset

We analyzed data from the Human Connectome Project Young Adult (HCP-YA) cohort (Van Essen et al., 2013; https://www.humanconnectome.org/, 2025 release). Each participant underwent a total of four 14.4-minute functional resting-state scans across two sessions, with phase-encoding directions applied alternately (1,200 frames per scan, repetition time [TR] = 0.72 s). The two sessions were acquired on separate days, with two scans per day. Throughout this article, session refers to one day of acquisition: we concatenated the two scans of a day into a single time series of 2,400 frames (about 29 minutes). Our analysis additionally incorporated the structural data of each participant, specifically the MSMAll-aligned midthickness surfaces. These surfaces provide the geodesic distances required for the surface-adapted clustering methods and the surface geometry required for the gradient-based method. We included 947 participants who completed all four resting-state scans, thereby holding scan duration constant across participants (Birn et al., 2013; Gordon et al., 2017) and retaining two independent sessions for the test-retest analyses. From these individuals, we also extracted cognitive measures and family relationships. Because the sample included twin and non-twin siblings, the 947 individuals were grouped into 419 distinct families.

### 2.2. Preprocessing

The 2025 HCP release provides resting-state data that have already undergone the minimal preprocessing pipelines (Glasser et al., 2013) followed by three noise-reduction stages: spatial independent component analysis (ICA) with FIX, in which components classified as noise were removed from the data (Salimi-Khorshidi et al., 2014), the Reclean procedure that refines FIX-based noise classification and removal (Yang et al., 2024), and temporal ICA that removes globally structured artifact while preserving the global signal (Glasser et al., 2018). The data have been surface-aligned using MSMAll (Robinson et al., 2014) and are provided in fs_LR 32k grayordinate space (Glasser et al., 2013; Van Essen et al., 2012).

To evaluate the robustness of our findings across different analytical decisions, we evaluated the data under four different preprocessing pipelines. Pipeline A used the HCP-provided data described above with no additional cleaning. Pipeline B expanded on this by applying nuisance regression, specifically detrending and the removal of 12 motion parameters, alongside a 0.01–0.10 Hz bandpass filter. Pipeline C applied global signal regression (GSR) directly to the minimally preprocessed data. Finally, Pipeline D incorporated all the nuisance regression and filtering steps from Pipeline B along with GSR. To avoid statistical artifacts, whenever multiple regressors were used (such as motion parameters and the global signal in Pipeline D), we regressed them out in a single step rather than sequentially. As a final step, we z-scored each vertex time series in all four pipelines. For Atlas-Guided Parcellation (AGP), Simple Linear Iterative Clustering (SLIC), and the gradient-based method, parcellations were estimated separately from the data produced by each preprocessing pipeline. The gradient-infused Multi-Session Hierarchical Bayesian Model (gMSHBM) parcellations were precomputed and therefore remained fixed across pipelines; only downstream time-series-derived measures were recomputed using each pipeline (Section 2.3.2).

### 2.3. Parcellation Methods

To ensure comparability and avoid dimensionality differences in the predictive models, we required all methods to produce exactly 200 parcels. The personalized parcellation methods differ in how they use group-level and individual data.

#### 2.3.1. Schaefer Group Atlas

Schaefer’s atlas, consisting of 200 parcels and 17 networks, was applied to each participant (Schaefer et al., 2018). The atlas was derived from a gradient-weighted Markov random field model that simultaneously optimized two criteria across 1,489 participants: local gradients in the connectivity profile, indicating where connectivity changes abruptly across the surface, and global similarity, requiring that vertices within a parcel share a similar whole-brain connectivity profile (Schaefer et al., 2018). Since the atlas applies the same boundaries across individuals, it serves as a group-level reference. Of the 59,412 cortical vertices in the data, 58,609 vertices fall within an atlas parcel; the remainder extend along the medial wall boundary and were left unassigned by all methods for comparability.

#### 2.3.2. Gradient-Infused Multi-Session Hierarchical Bayesian Model (gMSHBM)

The MS-HBM framework (Kong et al., 2019, 2021) estimates individual-specific brain networks by mathematically balancing group-level templates with the functional connectivity of each individual. This balance is governed by a hierarchical Bayesian model defined by distinct prior distributions. The group-level resting-state functional connectivity profile specifies the expected connectivity pattern of each parcel. The inter-individual prior models the expected spatial variation across the population, whereas the intra-individual prior accounts for scan-to-scan measurement noise within the same person. To prevent networks from scattering randomly across the cortex, the model applies a spatial localization prior. We used the gradient-infused variant (gMSHBM), which integrates local functional connectivity gradients into this prior. Rather than forcing strict spatial contiguity, this variant explicitly permits a single parcel to split into multiple non-contiguous components.

To ensure methodological replication, we leveraged the pre-computed individual gMSHBM parcellations directly from the authors’ repository (https://github.com/ThomasYeoLab/CBIG; Kong et al., 2021). Because we used these pre-calculated maps, the resulting parcellation boundaries reflect the preprocessing used in their original estimation. Consequently, unlike the other parcellation algorithms in our study, gMSHBM parcellations were not re-estimated under each of our four preprocessing pipelines. The same parcellation boundaries were therefore used across pipelines, while downstream time-series-derived measures were recomputed from the corresponding preprocessed data.

#### 2.3.3. Atlas-Guided Parcellation (AGP)

AGP takes only the seed positions from a group atlas, here the 200-parcel Schaefer atlas, and determines the extent and boundaries of every parcel using an iterative region-growing algorithm driven by the functional connectivity of each individual (Li et al., 2022). We initialized each parcel by placing a seed vertex at its geodesic center, defined as the vertex furthest from any boundary of the corresponding Schaefer parcel. Treating the medial wall as a boundary ensured these seeds remained well within the cortical sheet rather than near the edge. Because these coordinates depend solely on the static group atlas and surface mesh, we calculated them once and applied them uniformly across all participants. The parcels then grew outward from these seeds. At each step, the algorithm evaluated every unassigned vertex adjacent to a growing parcel and calculated its average correlation with the current members of each parcel:

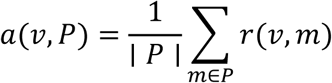

where *v* is a candidate vertex, *P* is the set of vertices currently assigned to that parcel, ∣ *P* ∣ is the number of vertices in that parcel, *m* is an individual vertex within *P*, and *r* is the Pearson correlation between their time series. The resulting value *a* represents the mean functional connectivity between the candidate vertex and the current members of the parcel. A global priority queue ranks these values, ensuring that the currently unassigned vertex–parcel pair with the highest mean functional connectivity is selected for assignment. Growth continued until the total number of assigned vertices matched the coverage of the group atlas.

#### 2.3.4. Simple Linear Iterative Clustering (SLIC)

SLIC (Wang et al., 2016) adapts the superpixel algorithm developed by Achanta et al. (2012) to parcellate the cortex. We adapted this framework for surface-based parcellation and implemented two distinct variants: a spatially constrained version (SLIC-F) and an unconstrained version (SLIC-C).

For the spatially constrained variant (SLIC-F), each vertex was assigned to the cluster center that minimized a combined functional and spatial distance metric (*d*):

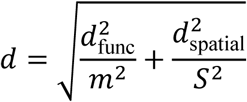

The spatial components of the equation encourage a spatial compactness constraint. Specifically, *d*_spatial_ is the geodesic distance, which measures the shortest path length along the folds of a structural midthickness mesh. 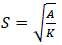 represents the expected spacing between parcels, where *K* is the number of parcels and *A* is the total physical surface area of the cortex. The functional component, *d*_func_, measures signal similarity. The original SLIC formulation defines this as the Euclidean distance between z-scored time series: 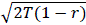, where *T* is the number of time points and *r* is the Pearson correlation. To decouple the metric from scan duration and ensure consistent mathematical scaling across both single– and dual-session data, we divided this term by √*T*, yielding a temporally normalized functional distance of 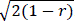. The compactness parameter *m* controls the relative weighting of the spatial and functional terms. We defined *m* as the median functional distance between adjacent vertices (estimated from 20 participants), using this value to scale the functional term relative to the spatial term in the SLIC distance metric. We used adjacent pairs rather than random pairs because distant cortical vertices are mostly uncorrelated; using random pairs would reduce the discriminative contribution of the functional term. Furthermore, because preprocessing steps (e.g., filtering) directly alter the functional distance scale, we re-estimated m separately for each of the four analytical pipelines to preserve comparable scaling between the functional and spatial terms.

For the unconstrained variant (SLIC-C), we removed the spatial constraint. SLIC-F therefore constrains individual functional clustering by a common spatial compactness criterion, whereas SLIC-C removes this spatial constraint and is driven solely by individual functional similarity. Vertices were clustered based solely on individual functional similarity using the equation: 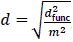. Consequently, SLIC-C allows parcels to form spatially disconnected components.

#### 2.3.5. Gradient-Based Parcellation

The gradient-based method (Gordon et al., 2016) delineates parcels by identifying locations on the cortical surface where functional connectivity patterns change abruptly. First, we defined the connectivity profile of every vertex by calculating a whole-cortex connectivity map, which is the vector of correlations between a given vertex *i* and all other cortical vertices. We then correlated these maps with one another to obtain the similarity between each pair of vertices:

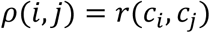

where *c_i_* is the connectivity map of vertex *i*. A high *ρ*(*i*, *j*) value indicates that the two vertices connect to the rest of the cortex in a similar way and therefore are more likely to belong to the same area.

In accordance with the reference implementation, the spatial gradient was calculated on a random subsample of these similarity maps (one map per 100 vertices, i.e., 594 out of 59,412 maps). This spatial gradient measures the local rate of change in connectivity across the surface mesh. The spatial gradient of each similarity map was then smoothed using a Gaussian kernel with *σ* = 2.55 mm. We performed these operations on each participant’s own MSMAll midthickness surface using Connectome Workbench (Marcus et al., 2011). Next, we applied the watershed-by-flooding algorithm to each gradient map (Vincent & Soille, 1991). This algorithm treats the gradient map as a topographical landscape: local minima (valleys of low change/high similarity) represent the centers of functional areas, and local maxima (ridges of high change) represent the boundaries. The proportion of maps in which a vertex coincided with a watershed boundary formed a final boundary probability map. We grew parcels outward from the local minimum points (the areas of lowest boundary probability) of this final boundary map. As the parcels grew, we merged neighboring parcels separated by weak boundaries, absorbed small parcels, and left vertices on strong boundaries unassigned as transition zones. Because each parcel grew directly from a local minimum across adjacent vertices of the surface mesh, and parcels were only merged with immediate neighbors, the method produces spatially connected parcels.

Unlike other approaches, this method does not constrain the final number of parcels. Our implementation yielded an average of 603 ± 22 parcels per participant, close to the 620.8 ± 39.4 parcels that the original authors reported for a different dataset, the Midnight Scan Club (Gordon et al., 2017). This similarity provides a numerical check on our implementation, although the datasets and the computational environment differ. To ensure comparability across methods, we required all methods to produce exactly 200 parcels. Because this specific algorithm stops merging when no boundary falls below a set threshold, we modified its default stopping criterion. We iteratively merged adjacent parcels sharing the weakest remaining boundary until exactly 200 parcels remained. We used these resolution-matched maps for all primary comparisons, while detailing the native-resolution outputs in Supplementary Material B.

#### 2.3.6. Hungarian Alignment

To enable cross-individual comparisons, we matched each individual parcel to the standard Schaefer atlas. We first quantified spatial agreement by constructing an overlap matrix *O*, where *O*(*p*, *a*) counts the vertices shared between an individual parcel *p* and an atlas parcel *a*. We then applied the Hungarian algorithm (Kuhn, 1955) to find the optimal one-to-one mapping *π* that solves:

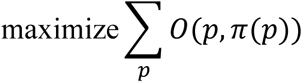

where *π*(*p*) represents the assigned atlas parcel for an individual parcel *p*. This mapping relabeled the individual parcels so that their total vertex overlap with the group atlas was maximal.

### 2.4. Validation Metrics

We evaluated four intrinsic properties of the personalized maps. These metrics characterize different aspects of the resulting parcellations and their downstream stability. Methods might achieve higher functional homogeneity at the cost of spatial coherence, or prioritize reliability by constraining individual variation.

#### 2.4.1. Functional Homogeneity

Homogeneity measures the functional similarity of the vertices within a parcel. For a parcel *p* containing vertices *V_p_*, with ∣ *V_p_* ∣ denoting the number of vertices, and having a mean time series of *t*ˉ*_p_*, homogeneity is the average correlation between each vertex and the parcel mean:

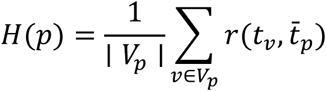

The reported value was the average of the *H*(*p*) values across all parcels. A higher score indicates higher functional similarity across vertices within a parcel.

Any method that explicitly maximizes within-parcel similarity, such as SLIC, can yield optimistically biased homogeneity estimates when homogeneity is evaluated on the same data used to derive the personalized parcellation. To avoid this circular evaluation, we assessed homogeneity out of sample by deriving the parcellation from one session and evaluating it on the other session. We excluded gMSHBM from this analysis because its parcellations were precomputed by the original authors using data combined from both sessions.

#### 2.4.2. Test-Retest Reliability

For AGP, SLIC-F, SLIC-C, and the gradient-based method, we independently estimated the parcellation from each session and used the corresponding session-specific map to extract the parcel-to-parcel functional connectivity (FC) matrix. To quantify test-retest reliability, we correlated the upper triangular elements of the two session-specific FC matrices:

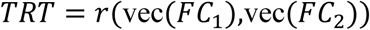

where vec(*FC_s_*) represents the vector of upper-triangular elements of the connectivity matrix obtained from session *s*. For AGP, SLIC-F, SLIC-C, and the gradient-based method, parcellations were estimated independently for each session, and the corresponding session-specific parcellation was used to derive the FC matrix. Thus, this metric captures the test-retest reproducibility of functional connectivity when both parcel boundaries and connectivity are re-estimated independently across sessions. For the methods that we applied to each session separately (Section 2.4.1), we also assessed the stability of the parcellation itself as the Dice overlap between a participant’s two single-session parcellations. This within-individual Dice captures how reproducible a participant’s own map is across the two days, and is therefore distinct from the inter-individual Dice coefficient (Section 2.4.4), which compares maps between different participants. For gMSHBM, session-specific parcellations were not available because the precomputed maps were derived from both sessions together; the same gMSHBM parcellation was therefore used to extract FC from each session, and its reliability estimate reflects connectivity stability under fixed parcel boundaries.

#### 2.4.3. Spatial Contiguity and Parcel Fragmentation

We defined a parcel as contiguous if all its vertices formed a single unfragmented component on the cortical surface. Two vertices were considered adjacent if they were connected by an edge in the cortical surface mesh. We calculated the percentage of contiguous parcels and the average number of disconnected components per parcel.

The Schaefer atlas and AGP guarantee contiguity by design, since the atlas parcels are contiguous by construction and AGP grows each parcel outward from a single seed. In the remaining methods parcels can fragment, and the fragments themselves may capture spatially disconnected topographic variation analogous to the variants described at the functional-network level (Dworetsky et al., 2024). To quantify this fragmentation, we identified the largest component of each fragmented parcel as its main body and calculated the shortest geodesic distance from each secondary fragment to this main body. This distance was used as a spatial descriptor of how far a secondary fragment was displaced from the main parcel component, rather than as a measure of functional impact. We excluded fragments containing fewer than 10 vertices to minimize spatial unreliability.

Our distance definition differs from the primary operationalization used by Dworetsky et al. (2024), who measured the distance of a network variant from the corresponding same-network boundary in a group-average map. Because our analysis focused on fragmentation of individual parcels rather than large-scale networks, we instead quantified displacement relative to the main body of the same personalized parcel. In our analysis, distance was therefore measured within each personalized parcel, from a secondary fragment to the main body of that same parcel. Accordingly, the 3.5 mm threshold reported by Dworetsky et al. was used only as a reference threshold rather than as an equivalent classification criterion. We evaluated this 3.5 mm value together with additional candidate thresholds (10, 15, and 30 mm) to examine how threshold choice affected the classification of candidate ectopic variants.

Rather than applying a single distance threshold to force a binary classification of fragments (i.e., border shift vs. ectopic variant), we report the full continuous distribution of these fragment distances. This approach explicitly allows us to evaluate how different distance thresholds change the detection of ectopic variants, accommodating potential variations introduced by data quality and spatial smoothing (Alakörkkö et al., 2017).

#### 2.4.4. Inter-Individual Dice Coefficient

The Dice coefficient quantitatively measures the degree of agreement between two participants’ parcellations. A lower number thus indicates a lower cross-individual spatial agreement. We measured the agreement through the shared assignment of vertex pairs:

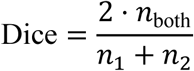

where *n_1_* and *n_2_* are the numbers of vertex pairs assigned to the same parcel in the first and the second participant, respectively, and *n_both_* is the number of pairs assigned to the same parcel in both participants.

### 2.5. Brain-Behavior Associations

#### 2.5.1. Neuroimaging Features

For the brain-behavior analyses, we used each participant’s parcellation derived from the concatenated resting-state data across both sessions. We extracted the mean time series of every parcel and computed four features for the brain-behavior analyses.

**(i)** **Global efficiency** A connectivity matrix was calculated at the parcel level and thresholded in three different ways: a weighted graph preserving all positive correlations, a binary graph preserving the strongest 10% of positive edges, and a binary graph preserving edges with *r* > .5. For each, global efficiency, that is, the average inverse shortest path length, was calculated as follows:

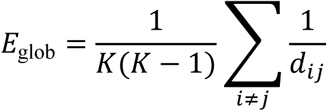

where *d_ij_* is the shortest path length between parcels *i* and *j*, and *K* = 200 is the number of parcels. Global efficiency indicates the potential for efficient communication or information transfer across the network and has been linked to individual differences in cognitive abilities (van den Heuvel et al., 2009).
**(ii)** **Intra-individual variability in parcel size** We computed the coefficient of variation (CV) of the 200 parcel sizes within each participant:

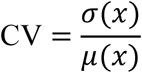

where *x* is the vector of the 200 parcel sizes for an individual, measured in vertices. The term *σ*(*x*) represents the standard deviation of those sizes, and *μ*(*x*) represents their mean. This scaling allows comparison of how unevenly the different methods partition the cortex while accounting for differences in mean parcel size. Individual differences in the spatial extent of functional regions have been shown to carry behavioral information, potentially reflecting meaningful variation in the spatial topography of functional networks across individuals (Bijsterbosch et al., 2018; Kong et al., 2019).
**(iii)** **Inter-individual parcel size similarity.** We calculated the mean correlation between the parcel size vector of each participant and those of all other participants. To ensure valid cross-individual comparisons, we ordered the elements of these vectors to match the standard Schaefer atlas indexing (via the alignment described in Section 2.3.6). Thus, parcel correspondence across individuals was defined relative to this common atlas-based labeling scheme. While the coefficient of variation quantifies size differences within a single brain, this metric evaluates how closely an individual’s pattern of parcel sizes resembles that of other participants. Consequently, individuals with less typical parcel-size profiles yield lower scores.
**(iv)** **Inter-individual connectivity similarity.** We calculated the mean correlation between the functional connectivity matrix of each participant and those of all other participants, also ordering the matrix rows and columns to match the standard atlas indexing. As for parcel size similarity, these comparisons therefore rely on the Schaefer-based correspondence established in Section 2.3.6. Because individual connectivity patterns are stable enough to act as unique identifiers across sessions (Finn et al., 2015), this measure of network typicality is a candidate feature for examining brain-behavior associations.

Features (ii) and (iii) exist only for personalized parcellations because the static group atlas assigns the exact same parcel sizes to every participant. We therefore excluded these two spatial metrics from the analyses of the group atlas.

#### 2.5.2. Cognitive Performance

We estimated general intelligence (*g*) using nine tasks from the HCP test battery. We modeled this construct using the established bifactor structure proposed by Dubois et al. (2018). Specifically, we specified a confirmatory factor analysis (CFA) model where a single general factor, representing *g*, loaded on all nine cognitive tasks. To capture domain-specific variance independent of *g*, we included three orthogonal group factors: processing speed (Dimensional Change Card Sort, Flanker, Pattern Completion Processing Speed), crystallized ability (Picture Vocabulary, Oral Reading Recognition), and memory (Penn Word Memory, Picture Sequence Memory). Following the pre-defined architecture of Dubois et al. (2018), the Penn Progressive Matrices and Variable Short Penn Line Orientation tests loaded only on the general factor. Because the crystallized ability and memory factors each contain only two indicators, their loadings cannot both be freely estimated while uniquely identifying these specific factors. We therefore followed standard identification practice and constrained the two loadings within each of these factors to be equal. We standardized all test scores prior to model estimation. We excluded seven participants entirely from this model because they lacked valid scores for one or more of the nine required cognitive tasks. This exclusion left a final sample of 940 participants for all behavioral analyses.

#### 2.5.3. Brain-Behavior Modeling

**(i) Structural equation modeling (SEM) with fixed loadings.** We used a two-step approach to examine the associations between brain features and g. In the first step, we estimated a measurement model in which *g* was defined by the nine cognitive tasks, as described in Section 2.5.2. We then extracted the unstandardized factor loadings, which quantify how strongly each cognitive task is related to *g*. In the second step, we introduced the brain features as predictors of *g*, while keeping the factor loadings fixed at their previously estimated values. This approach was intended to ensure that the definition of *g* remained consistent across analyses. If the factor loadings and the regression paths from brain features to *g* were estimated simultaneously, the factor loadings could change depending on the brain features included in the model. Consequently, different sets of brain features could be associated with slightly different definitions of *g*, making their associations difficult to compare. By fixing the factor loadings, we ensured that all brain features were evaluated against the same behaviorally defined latent construct, rather than allowing its measurement to adapt to the brain features being tested.

To evaluate how different aspects of network architecture are associated with intelligence, we tested combinations of brain features. Every model included inter-individual connectivity similarity. We then varied the operationalization of global efficiency (using three different graph thresholds to account for both dense and sparse network topologies) and spatial topography (using three different parcel size configurations). For each personalized parcellation method, this approach yielded nine distinct structural equation models (Table 1). Because the static group atlas (Schaefer) assigns identical parcel sizes to every participant, it lacks spatial variance. We therefore evaluated the group atlas using only Models 1 through 3.

**Table 1.** Configuration of the nine structural equation models. CV represents the coefficient of variation of the 200 parcel sizes within each participant

| Model | Global Efficiency Threshold | Spatial Topography Feature | Constant Feature (All Models) |
| --- | --- | --- | --- |
| 1 | Weighted graph (all positive $r$) | None | Connectivity similarity |
| 2 | Binary graph (top 10% edges) | None | Connectivity similarity |
| 3 | Binary graph ($r > .5$) | None | Connectivity similarity |
| 4 | Weighted graph (all positive $r$) | CV of parcel sizes | Connectivity similarity |
| 5 | Binary graph (top 10% edges) | CV of parcel sizes | Connectivity similarity |
| 6 | Binary graph ($r > .5$) | CV of parcel sizes | Connectivity similarity |
| 7 | Weighted graph (all positive $r$) | Parcel size similarity | Connectivity similarity |
| 8 | Binary graph (top 10% edges) | Parcel size similarity | Connectivity similarity |
| 9 | Binary graph ($r > .5$) | Parcel size similarity | Connectivity similarity |

Standardized regression paths report the association between each predictor and g while holding all other variables constant. This approach can reduce the estimated contribution of predictors that share variance with one another. To quantify the unique variance associated with each brain feature, we refit each model, removing one predictor at a time. We defined the resulting drop in explained variance as its unique contribution:

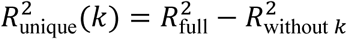

where *k* denotes the predictor removed from the model.

**(ii) Kernel ridge regression.** While our SEM analysis evaluates the associations of selected summary features with *g*, we used kernel ridge regression (KRR) as a multivariate approach to test how well *g* can be predicted from the high-dimensional data generated by a parcellation (Kong et al., 2019, 2021). For each personalized parcellation method, we derived two distinct prediction kernels: one capturing participant similarity based on the full vector of parcel sizes, and another capturing participant similarity based on the full functional connectivity matrix. These kernels represent complementary information carried by the parcellations: spatial topography through parcel-size variation and functional organization through inter-regional connectivity, both of which have previously been linked to individual differences in behavior (Kong et al., 2019, 2021). For the Schaefer atlas, only the connectivity kernel was available because parcel sizes do not vary across participants. Prior to prediction, we regressed out age and sex, and centered the kernel strictly within the training folds to prevent data leakage. We split the data into 10 cross-validation folds based on family structure, ensuring that related individuals remained grouped together in either the training or test set. Within each fold, we selected the optimal ridge regularization parameter using the closed-form leave-one-out cross-validation error (Kong et al., 2019). We quantified prediction accuracy as the Pearson correlation between the observed and predicted *g* scores. To assess statistical significance, we shuffled the behavioral outcomes 1,000 times to create an empirical null distribution and compared our observed accuracy against it. Finally, to account for multiple testing across all combinations of parcellation methods and kernels, we adjusted the empirical *p*-values to control the false discovery rate (FDR).

### 2.6. Statistical Analysis

Because we applied all six parcellation approaches to the same set of participants, we evaluated performance differences using within-person statistical designs. For metrics where all methods exhibited variance, we tested for differences using Greenhouse-Geisser-corrected repeated-measures ANOVAs, reporting generalized eta-squared (*η*^2^) as the effect size. For contiguity and inter-individual Dice, some methods produced constant values across participants by construction. For example, the Schaefer atlas has fixed contiguity and inter-individual Dice values, while AGP always yields 100% contiguity because parcel growth is restricted to adjacent surface vertices. We therefore compared these metrics across all methods using the rank-based Friedman test, which accommodates such tied or constant values, and reported Kendall’s *W* as the effect size. We evaluated pairwise differences between methods using Holm-corrected paired t-tests and reported Cohen’s *d* (Supplementary Material E). Pairwise tests were not performed when two methods produced identical values for every participant.

We estimated all structural equation models, including the confirmatory factor analysis for cognitive performance described in Section 2.5.2, using robust maximum likelihood. To account for the family structure in the HCP dataset, we clustered standard errors at the family level. We evaluated model fit using the robust comparative fit index (CFI), Tucker-Lewis index (TLI), root mean square error of approximation (RMSEA), and standardized root mean square residual (SRMR). Because we fit multiple structural models for each preprocessing pipeline, we corrected the *p*-values for multiple comparisons. Specifically, for a given pipeline, we pooled the *p*-values from all structural regression paths and adjusted them using the false discovery rate (FDR).

### 2.7. Software Implementation

We performed the analyses in `Python` 3.9 and `R` 4.3.1. Within `Python`, we used `nibabel` for CIFTI surface operations, `NumPy` and `SciPy` for numerical computation, and `nilearn` for visualization. We executed the spatial gradient and geodesic smoothing steps using `Connectome Workbench` 1.5.0, following the gradient-based method’s reference implementation. We ran all statistical analyses in `R`, using `lavaan` 0.6-15 for structural equation modeling and `afex` for the repeated-measures tests.

The original parcellation algorithms span multiple programming languages and environments. We reimplemented the three algorithms that we ran ourselves (AGP, SLIC, and the gradient-based method) into a single Python codebase with a unified interface to standardize the inputs and outputs. We validated each reimplementation against its original formulation (see numerical checks in Supplementary Material C). The complete analysis pipeline is available at https://github.com/CoskuInceler/Individualized_Brain_Parcellation; access to the HCP data requires separate authorization.

### 2.8. Statement of the Use of AI Tools

Google Gemini and OpenAI ChatGPT were used to assist with language editing and refinement of the manuscript. Anthropic Claude Code was used to assist with code development, debugging, and refinement. All AI-assisted text and code were critically reviewed and verified by the authors, who take full responsibility for the content of the manuscript, the analysis code, and the reported results.

## 3. Results

Sections 3.1 through 3.4 present the results obtained from the minimal preprocessing pipeline (pipeline A). Section 3.5 examines whether these findings hold for the remaining three pipelines (pipelines B, C, and D).

### 3.1. Spatial Topography Separates Parcellation Methods

First, we evaluated the spatial and geometric characteristics of the parcels generated by each method. Figure 1 illustrates the cortical maps obtained for three representative participants. While the group atlas remains identical across all participants by construction, the five personalized methods generated visibly distinct boundaries for each individual. Beyond these individual differences, the algorithms diverged in how they shaped the parcels. AGP, SLIC-F, and the gradient-based method yielded compact, spatially contiguous areas. In contrast, gMSHBM and SLIC-C produced highly fragmented parcels.

**Figure 1.**
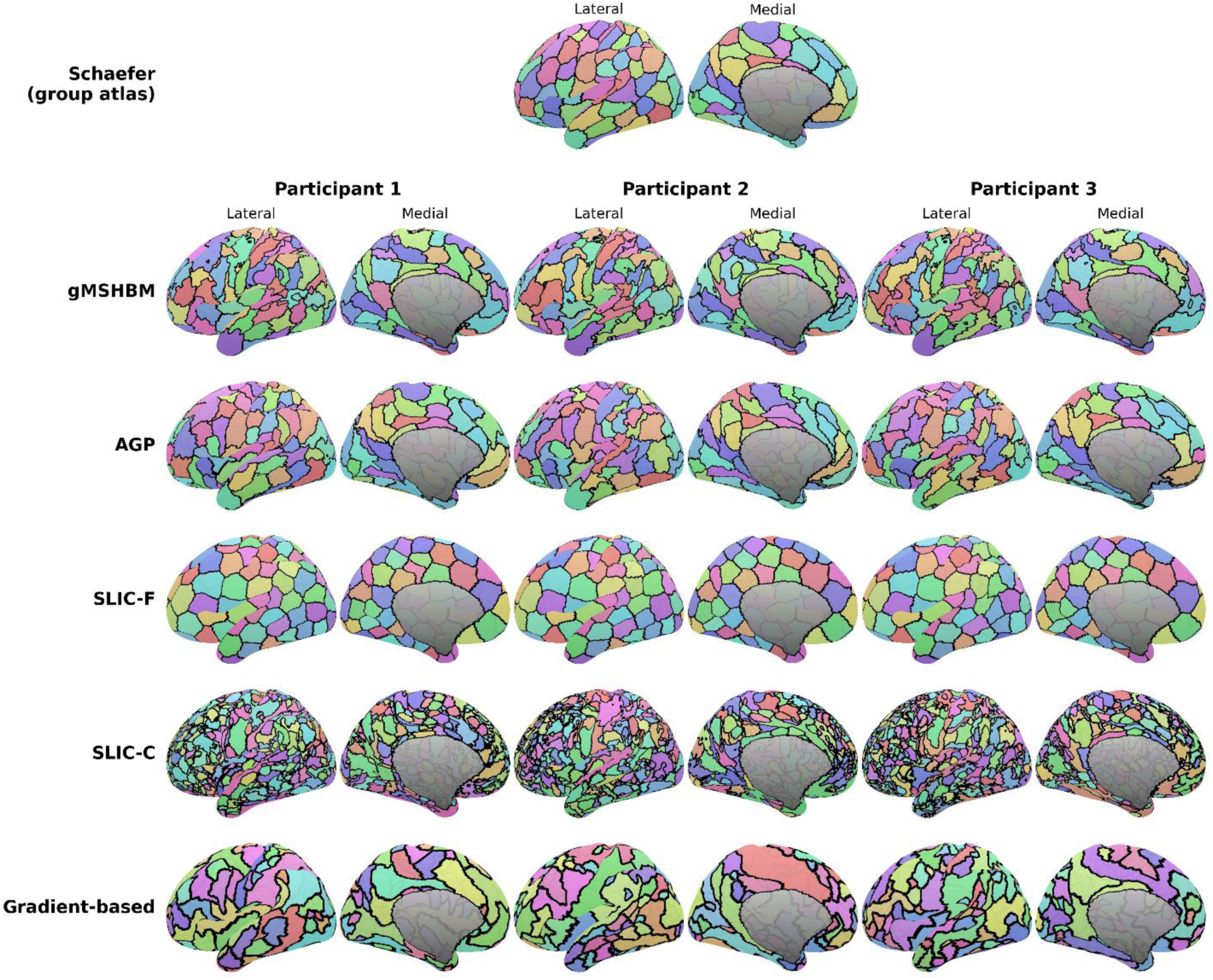
Parcellations produced by each method (left hemisphere, lateral and medial views). The Schaefer group atlas is shown only once, as it remains identical across all participants. We display the five personalized methods (rows) for the same three individuals (columns). All outputs were standardized to a 200-parcel resolution (the gradient-based method was iteratively merged down from a native resolution of ∼600). The algorithms yield visibly distinct topographies: AGP, SLIC-F, and the gradient-based method generate spatially contiguous parcels, whereas gMSHBM and SLIC-C produce fragmented parcels comprising multiple disconnected components. Abbreviations: AGP, Atlas-Guided Parcellation; gMSHBM, gradient-infused Multi-Session Hierarchical Bayesian Model; SLIC-C, Simple Linear Iterative Clustering (unconstrained); SLIC-F, Simple Linear Iterative Clustering (spatially constrained).

We evaluated the spatial and functional properties of the parcels (Figure 2). Functional homogeneity (Figure 2A) differed across methods (Greenhouse-Geisser corrected *F*(2.33, 2204) = 31,110.6, *p* < .001, generalized *η*² = .51). The gradient-based method achieved the highest value (.379), followed by SLIC-C (.373). The remaining four methods clustered within a narrow range: gMSHBM .318, AGP .316, Schaefer atlas .309, and SLIC-F .305. Post-hoc Holm-corrected paired *t*-tests confirmed that all pairwise differences between methods were statistically significant (all *p* < .001), although the magnitude of these within-participant differences varied substantially across method pairs (*d*_av_ = 0.07 to 2.53; Supplementary Material E).

**Figure 2.**
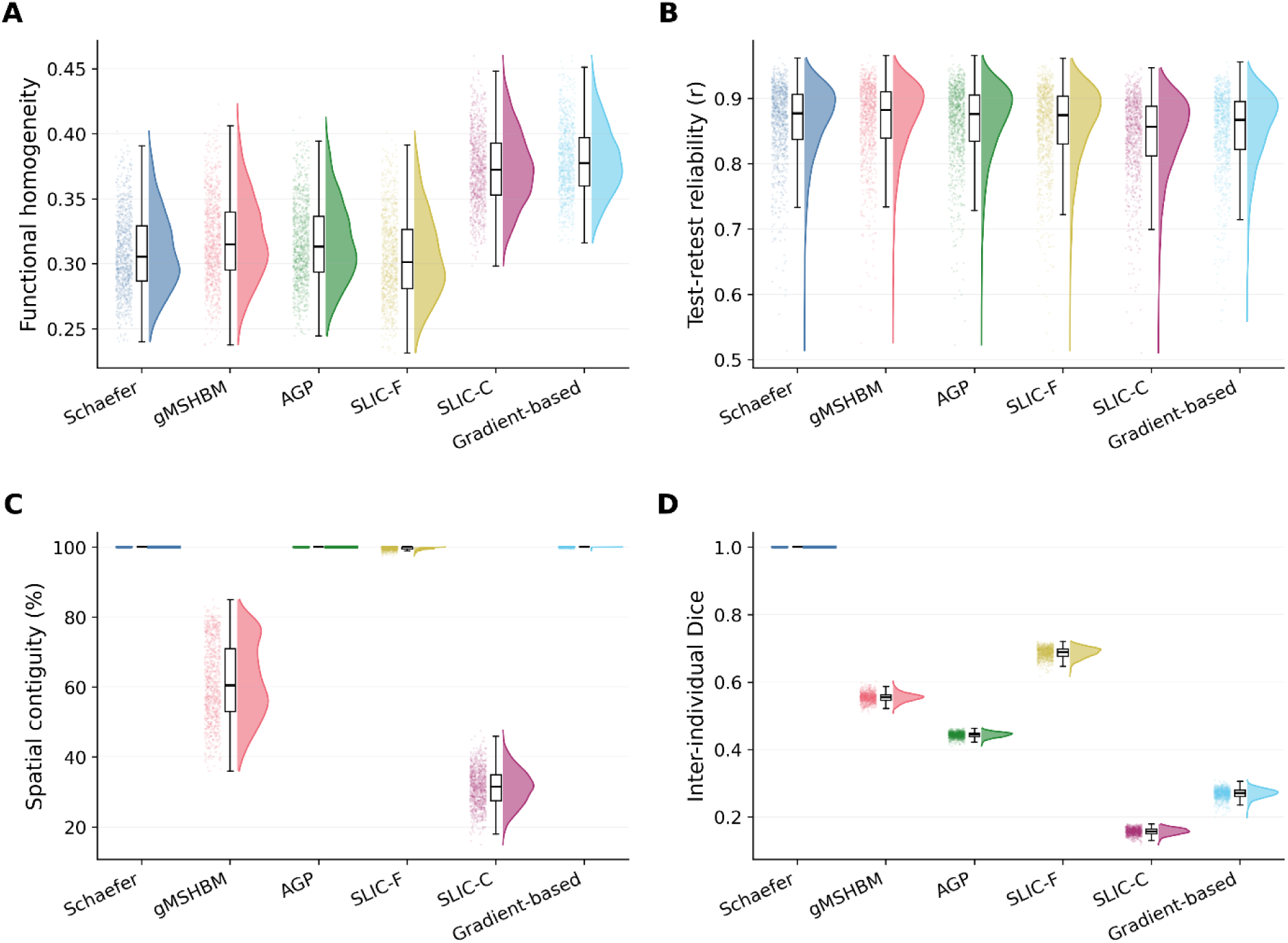
Validation metrics across the six methods. (A) functional homogeneity, (B) test-retest reliability of connectivity, (C) spatial contiguity, and (D) inter-individual Dice coefficient. Panels show data distributions (half-violins), medians and quartiles (boxes), and individual points. Functional homogeneity was highest for SLIC-C and the gradient-based method, whereas test-retest reliability of connectivity was similar across methods. Spatial contiguity was highest for Schaefer, AGP, SLIC-F, and the gradient-based method, but substantially lower for gMSHBM and SLIC-C. Inter-individual Dice showed pronounced differences across all personalized methods. The Schaefer atlas (contiguity and Dice) and AGP (contiguity) appear as single lines rather than distributions because these metrics are fixed at their maximum values by construction, resulting in zero variance. Abbreviations: AGP, Atlas-Guided Parcellation; gMSHBM, gradient-infused Multi-Session Hierarchical Bayesian Model; SLIC-C, Simple Linear Iterative Clustering (unconstrained); SLIC-F, Simple Linear Iterative Clustering (spatially constrained).

Test-retest reliability of connectivity (Figure 2B) showed the least variation among the four metrics. All six methods fell within the narrow range of .842 to .867, and method choice explained less than 2% of the variance (*F*(2.35, 2223) = 839.3, *p* < .001, generalized *η*² = .018). gMSHBM yielded the highest reliability (.867), followed by the Schaefer atlas (.863), AGP (.862), SLIC-F (.859), the gradient-based method (.852), and SLIC-C (.842). All pairwise differences were significant after Holm correction (all *p* < .001), although the magnitude of these differences was small (*d*_av_ = 0.02 to 0.41).

Spatial contiguity (Figure 2C) divided the methods into two groups (Friedman, *χ*²(5) = 4,455.9, *p* < .001, *W* = .94). Contiguity for the Schaefer atlas and AGP was exactly 100%, as expected given the algorithmic constraints. The gradient-based method reached 99.9%, and SLIC-F reached 99.6%. In contrast, contiguity dropped to 61.5% for gMSHBM and 31.2% for SLIC-C. All pairwise differences were significant after Holm correction (all *p* < .001; *d*_av_ = 0.32 to 26.16). The Schaefer atlas and AGP could not be compared, since both were at exactly 100% in every participant.

The inter-individual Dice coefficient (Figure 2D) ranked the methods consistently across the entire sample (Friedman, *χ*²(5) = 4,735.0, *p* < .001, *W* = 1.00, meaning all 947 participants produced the identical ranking). The Schaefer atlas scored 1.00 because it has no variability, which anchors this perfect rank consistency. Among the personalized methods, SLIC-F showed the highest overlap (.687), followed by gMSHBM (.554), AGP (.443), the gradient-based method (.269), and SLIC-C (.157). All pairwise differences were significant after Holm correction (all *p* < .001), with uniformly large effect sizes (*d*_av_ = 9.0 to 169.4).

Evaluating homogeneity on the same data used to estimate the parcellation introduces circularity. We therefore calculated homogeneity across independent sessions to assess out-of-sample performance. The difference between within-session and cross-session values was .002 for SLIC-F, .002 for the gradient-based method, .003 for AGP, and .010 for SLIC-C. Thus, cross-session homogeneity was approximately 3% lower than within-session homogeneity for SLIC-C, whereas the differences were minimal for the other three methods.

Beyond the reliability of functional connectivity, we assessed the spatial stability of the parcellations. The within-individual Dice coefficient between a participant’s two single-session parcellations was .761 for SLIC-F, .592 for AGP, .318 for the gradient-based method, and .255 for SLIC-C. We excluded gMSHBM from this analysis because its parcellations were precomputed by the original authors using both sessions together.

### 3.2. Ectopic Variant Rates Are Method– and Threshold-Dependent

As established in Section 3.1, the methods differ in how frequently they fragment parcels. To evaluate whether secondary fragments would be classified as local border shifts or candidate ectopic variants, we measured the geodesic distance between each secondary fragment and the main body of its parcel.

The four methods varied substantially in both fragment count and spatial dispersion (Figure 3A). The gradient-based method produced almost no fragments (26 fragments across 25 participants). SLIC-F generated 692 fragments across 460 participants, nearly all located within a few millimeters of the main body. gMSHBM produced fragments for every participant (107,317 total); although mostly concentrated nearby, some extended much further away. SLIC-C produced substantially more fragments, generating 1,914,392 fragments distributed broadly across all distances.

**Figure 3.**
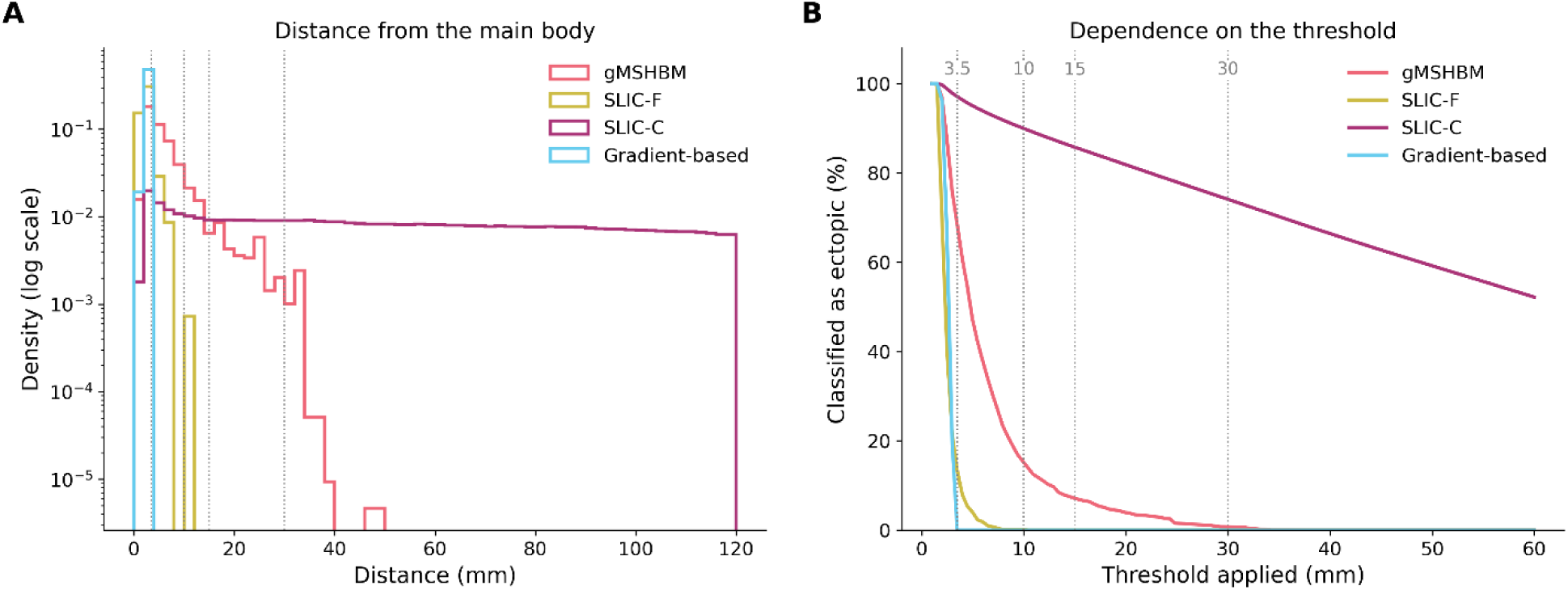
Distance of parcel fragments from the main body. (A) Distribution of distances for the four methods that produced fragments, plotted on a logarithmic density scale. Distances were measured from each secondary fragment to the main body of the corresponding personalized parcel, with larger distances indicating greater spatial separation between parcel components. Candidate thresholds are marked as dotted lines; distances exceeding 120 mm are omitted. (B) Proportion of fragments classified as candidate ectopic variants at each distance threshold. The 3.5 mm threshold was adopted as a reference from Dworetsky et al. (2024), although their primary classification measured distance relative to a group-average network map rather than an individual’s parcel main body. For gMSHBM, 68.2% of fragments are located beyond 3.5 mm but only 0.7% fall beyond 30 mm. In contrast, most SLIC-C fragments exceed every candidate threshold, consistent with SLIC-C allowing functionally similar vertices to be assigned to the same parcel without a spatial compactness constraint. The gradient-based method produced 26 fragments across 25 participants, all located within 3.5 mm of the main body. AGP produced no fragments and is therefore not shown. Abbreviations: AGP, Atlas-Guided Parcellation; gMSHBM, gradient-infused Multi-Session Hierarchical Bayesian Model; SLIC-C, Simple Linear Iterative Clustering (unconstrained); SLIC-F, Simple Linear Iterative Clustering (spatially constrained).

Consequently, the proportion of fragments classified as ectopic depended heavily on the specific distance threshold (Figure 3B). Applying the 3.5 mm threshold used by Dworetsky et al. (2024) at the network level, while measuring distance here from each fragment to the main body of the corresponding personalized parcel, 68.2% of gMSHBM fragments would be classified as candidate ectopic variants. This classification rate dropped to 15.2% at 10 mm, 7.1% at 15 mm, and just 0.7% at 30 mm. SLIC-C was less sensitive to threshold selection, with 97.0% of fragments exceeding 3.5 mm and 74.1% remaining beyond 30 mm. Among the rare SLIC-F fragments, 13.0% exceeded 3.5 mm, and almost none exceeded 10 mm. No gradient-based fragments exceeded the 3.5 mm threshold.

### 3.3. Network Topology Differs Between Constrained and Unconstrained Parcellations

We next evaluated how graph metrics varied across parcellation methods. Global efficiency differed across methods (*F*(1.33, 1261) = 4,033.2, *p* < .001, generalized *η*² = .28). Among the four constrained methods (Schaefer, gMSHBM, SLIC-F, and AGP), global efficiency for the weighted graph fell within a narrow range (Schaefer: .201; gMSHBM: .204; SLIC-F: .200; AGP: .194). In contrast, the two unconstrained methods yielded substantially lower efficiencies (SLIC-C: .161; gradient-based method: .162; Figure 4A). Differences among the four constrained methods were negligible (*d*_av_ = 0.02 to 0.31), as was the difference between the two unconstrained methods (*d*_av_ = 0.07), whereas every constrained-unconstrained comparison was large (*d*_av_ = 1.19 to 1.65).

**Figure 4.**
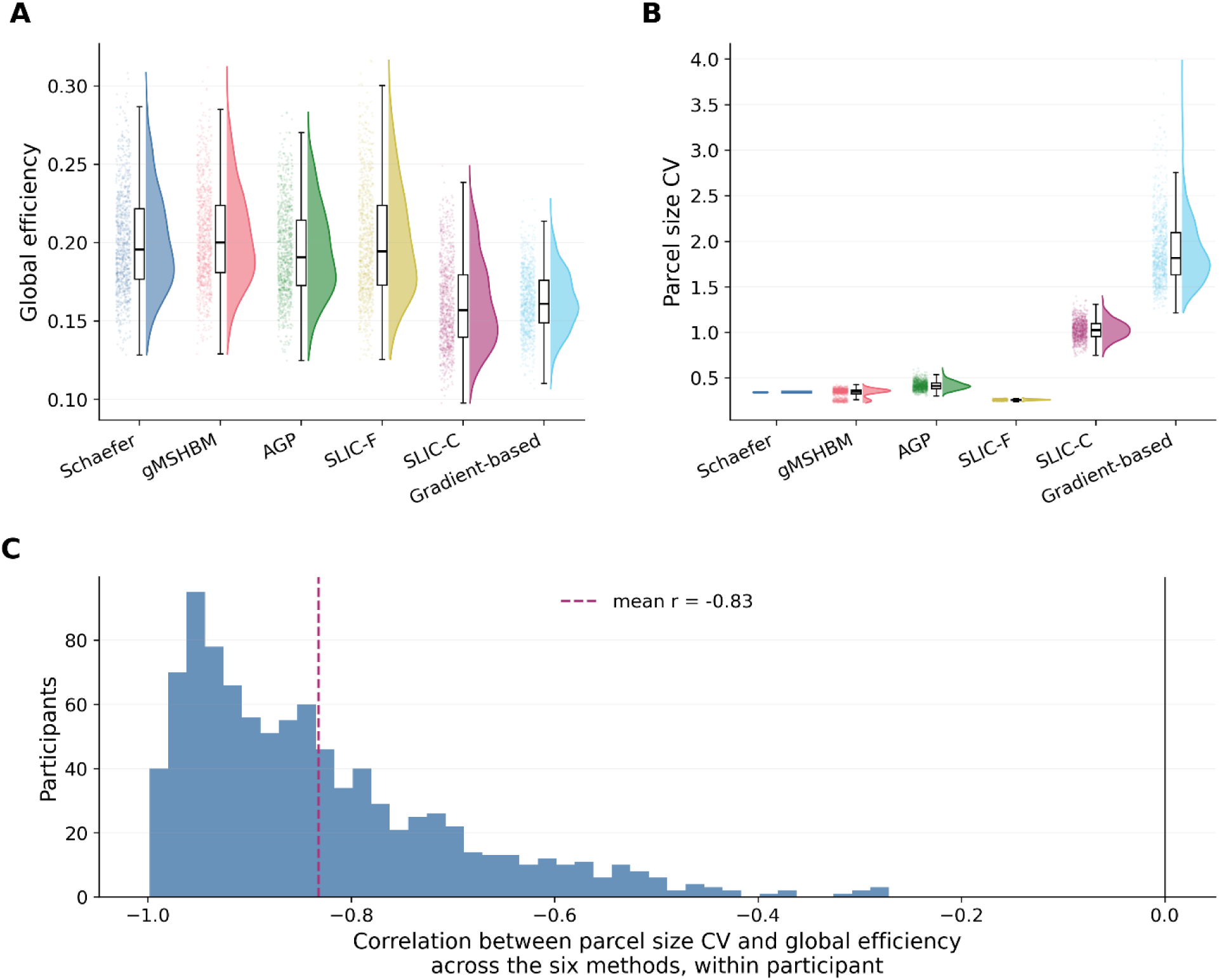
Graph-level properties of the parcellations. (A) Global efficiency of the weighted graph and (B) the coefficient of variation of parcel size, shown as data distributions (half-violins), medians and quartiles (boxes), and individual participants (points). The four constrained methods overlap in global efficiency and show lower parcel size variability, whereas SLIC-C and the gradient-based method show lower global efficiency and higher parcel size variability. The Schaefer atlas appears as a single line in (B) because its parcel sizes are fixed by the atlas. (C) Distribution of the within-participant correlation between parcel size CV and global efficiency computed across the six methods, one value per participant. The correlation is negative in every participant, showing that, across methods within individuals, greater parcel size variability is associated with lower global efficiency. Abbreviations: AGP, Atlas-Guided Parcellation; CV, coefficient of variation; SLIC-C, Simple Linear Iterative Clustering (unconstrained); SLIC-F, Simple Linear Iterative Clustering (spatially constrained).

This difference in global efficiency coincided with substantial differences in parcel size variability. The coefficient of variation for parcel size remained low for the constrained methods (SLIC-F: .260; gMSHBM: .337; Schaefer: .343; AGP: .417). However, the coefficients of variation exceeded 1 for both unconstrained methods (SLIC-C: 1.027; gradient-based: 1.890; Figure 4B). The gradient-based method divided the cortex particularly unevenly; in a representative participant, parcel sizes ranged from 19 to 4,036 vertices (median = 94). Within each participant, parcel size variability and global efficiency were strongly and negatively related across the six methods (mean *r* = −.83, *SD* = .13), and this relationship was negative in all 947 participants (Figure 4C).

Applying binary thresholds increased graph fragmentation. Retaining only the strongest 10% of positive edges fractured the 200-node graph into multiple disconnected components (Schaefer and gMSHBM: 35; AGP: 37; SLIC-F: 40; gradient-based: 47; SLIC-C: 55). Applying the stricter threshold (*r* > .5) split the graphs into even more isolated components. Because disconnected pairs of parcels contribute zero to the calculation, this increasing graph fragmentation contributed to the lower global efficiency observed under the stricter thresholds.

### 3.4. Topographical Variability Is Associated with Behavior Primarily Under gMSHBM

The bifactor measurement model fit the cognitive data well (CFI = .970, TLI = .950, RMSEA = .051 [.038, .065], SRMR = .032). All nine tasks loaded significantly on the general factor (Supplementary Material D). The Penn Progressive Matrices (.69), Oral Reading Recognition (.65), Picture Vocabulary (.61), and Line Orientation (.59) tasks yielded the highest loadings. The speed tasks yielded among the lowest loadings (.23 to .31), as their own group factor absorbed most of their variance. We fixed these loadings to isolate and estimate only the regression paths from the brain features to *g*.

Only six of the 126 regression paths evaluated across all thresholds survived correction for multiple comparisons. These six paths represent two spatial predictors under gMSHBM, both of which remained significant across all three graph thresholds (Figure 5A shows the weighted graph). Specifically, the coefficient of variation of gMSHBM parcel sizes was positively associated with *g* consistently across all three graph thresholds (*β* = .220, .220, and . 214), and gMSHBM parcel size similarity was likewise positively associated with *g* across all three thresholds (*β* = .192, .171, and . 200). No path from global efficiency, connectivity similarity, or any other parcellation method reached significance. Because these associations remained virtually identical across all thresholds, Table 2 reports the detailed model parameters for the weighted graph as representative estimates. The estimates for the two binary thresholds are reported in Supplementary Material F.

**Figure 5.**
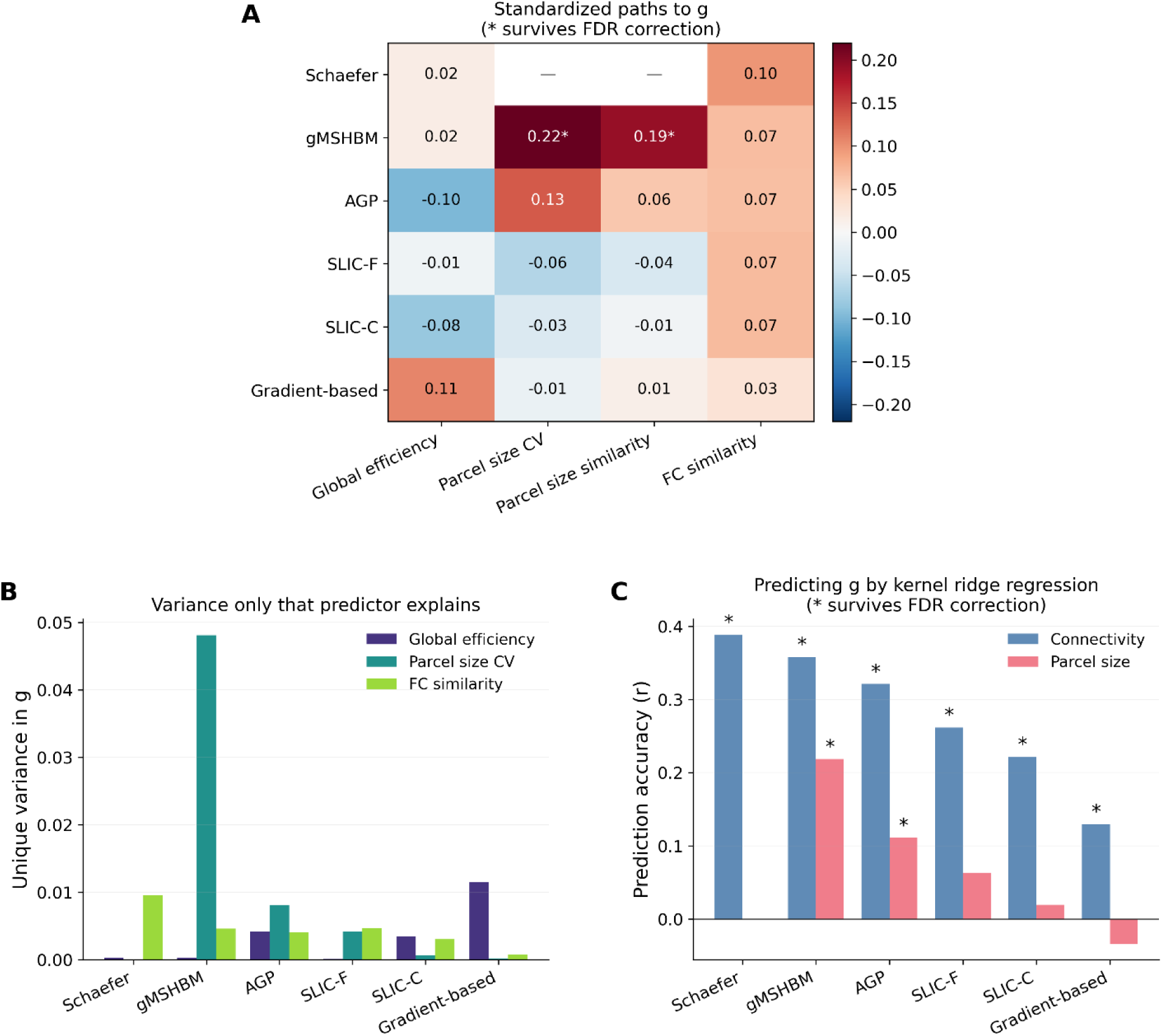
Brain-behavior associations and prediction of general intelligence (*g*) across the six parcellation methods. (A) Standardized regression paths from the structural equation models. Predictors include global efficiency, the coefficient of variation of parcel size, parcel size similarity, and functional connectivity similarity. The Schaefer group atlas lacks spatial variance due to algorithmic constraints, leaving the spatial predictor columns empty. Asterisks mark paths that survive false discovery rate correction. (B) Unique variance in *g* associated with individual predictors, quantified as the reduction in model R² when each predictor was removed. (C) Out-of-sample prediction accuracy (Pearson *r*) from the kernel ridge regression models. Predictions rely on either a functional connectivity kernel or a parcel size kernel. Asterisks denote prediction accuracies that survive false discovery rate correction. Abbreviations: CV, coefficient of variation; FC, functional connectivity; FDR, false discovery rate.

**Table 2.** Standardized regression paths from brain features to *g*. Estimates are shown for the weighted graph, with factor loadings fixed to the values obtained from the measurement model fitted to the cognitive tasks alone. Each row represents one method under one predictor set. The set labeled “CV” included the coefficient of variation of parcel sizes, the set labeled “similarity” included parcel size similarity, and the set labeled “none” included no parcel size feature. Global efficiency and connectivity similarity entered every set. Asterisks denote significance after false discovery rate correction across all 126 regression paths (\*\*\**p* < .001). Dashes mark predictors not included in that set or unavailable for that method. Only the two parcel-size predictors under gMSHBM survived correction. Abbreviations: AGP, Atlas-Guided Parcellation; CV, coefficient of variation; FC, functional connectivity; gMSHBM, gradient-infused Multi-Session Hierarchical Bayesian Model; SLIC-C, Simple Linear Iterative Clustering (unconstrained); SLIC-F, Simple Linear Iterative Clustering (spatially constrained).

| Method | Predictor set | FC similarity | Global efficiency | Parcel size CV | Parcel size similarity |
| --- | --- | --- | --- | --- | --- |
| Schaefer | none | .098 | .017 | — | — |
| gMSHBM | CV | .068 | .017 | **.220\*\*\*** | — |
| gMSHBM | similarity | .021 | .072 | — | **.192\*\*\*** |
| gMSHBM | none | .081 | .019 | — | — |
| AGP | CV | .067 | -.100 | .134 | — |
| AGP | similarity | .033 | .002 | — | .059 |
| AGP | none | .052 | .003 | — | — |
| SLIC-F | CV | .070 | -.010 | -.065 | — |
| SLIC-F | similarity | .082 | -.022 | — | -.035 |
| SLIC-F | none | .070 | -.010 | — | — |
| SLIC-C | CV | .069 | -.083 | -.032 | — |
| SLIC-C | similarity | .073 | -.066 | — | -.007 |
| SLIC-C | none | .072 | -.066 | — | — |
| Gradient-based | CV | .031 | .108 | -.014 | — |
| Gradient-based | similarity | .036 | .106 | — | .012 |
| Gradient-based | none | .037 | .107 | — | — |

Variance decomposition mirrored these path estimates (Figure 5B). The full gMSHBM model explained 5.5% of the variance in *g*, whereas the other five methods explained only 0.5% to 1.3%. Most of the variance explained by the gMSHBM model was attributable to the parcel size coefficient of variation. Removing the gMSHBM parcel size coefficient of variation dropped the explained variance by 4.8 percentage points. This single feature constituted 88% of the total variance explained by the model. The corresponding unique variance contributions for the other methods were negligible (AGP: .008; SLIC-F: .004; SLIC-C: .001; gradient-based: .0002).

While the structural equation models tested whether isolated summary metrics (e.g., the overall coefficient of variation) were associated with *g*, kernel ridge regression addresses a different question. It evaluates the out-of-sample predictive information contained in the entire topography, specifically the full vector of 200 parcel sizes or the complete 200 × 200 connectivity matrix (see Methods). Using the connectivity matrix kernel, the Schaefer atlas achieved the highest accuracy (*r* = .389), followed by gMSHBM (*r* = .358), AGP (*r* = .321), SLIC-F (*r* = .262), SLIC-C (*r* = .222), and the gradient-based method (*r* = .130; all FDR-corrected *p* < .01; Figure 5C).

Using the parcel size kernel, gMSHBM achieved the highest accuracy (*r* = .218, *p* < .01), followed by AGP (*r* = .111, *p* < .01). Predictions from SLIC-F (*r* = .063), SLIC-C (*r* = .019), and the gradient-based method (*r* = –.034) were not significant. The Schaefer atlas lacks parcel size variance across participants and was therefore excluded from this specific spatial analysis (Table 3).

**Table 3.** Prediction of *g* using KRR and variance explained by the SEM models. Kernel ridge regression accuracy for the two prediction kernels, with significance assessed by permutation against 1,000 shuffles of the outcome. Values are corrected using the false discovery rate (** FDR-corrected *p* < .01). Cross-validation folds were assigned by family rather than by participant. The SEM *R*^2^ column reports the total variance in *g* explained by the full structural equation model. The final column gives the unique contribution of parcel size variability, computed as the drop in explained variance when that predictor alone is removed. The two KRR kernels yielded different between-method patterns: the connectivity kernel showed the highest accuracy for the Schaefer atlas, whereas the parcel size kernel showed the highest accuracy for gMSHBM. The Schaefer atlas carries no parcel size variance and is therefore absent from both parcel size columns. Abbreviations: FDR, false discovery rate; KRR, kernel ridge regression; SEM, structural equation modeling.

| Method | KRR $r$ (connectivity) | KRR $r$ (parcel size) | SEM $R^2$ | Unique $R^2$ (parcel size) |
| --- | --- | --- | --- | --- |
| Schaefer | .389** | — | .0098 | — |
| gMSHBM | .358** | **.218\*\*** | **.0549** | **.0481** |
| AGP | .321** | .111** | .0109 | .0081 |
| SLIC-F | .262** | .063 | .0090 | .0042 |
| SLIC-C | .222** | .019 | .0045 | .0007 |
| Gradient-based | .130** | -.034 | .0125 | .0002 |

### 3.5. Preprocessing Choices Change the Number of Detected Brain-Behavior Associations

We reported the preceding results using minimally preprocessed data (pipeline A). To assess robustness, we repeated the analyses under three alternative preprocessing pipelines (Supplementary Material A). For gMSHBM, the precomputed parcellations were held fixed across pipelines, whereas the downstream time-series-derived features were recomputed. The core spatial metrics showed largely consistent between-method patterns. Across all four pipelines, the relative ranking of methods by inter-individual Dice coefficient remained identical. The sharp division in spatial contiguity also persisted.

The relative pattern of predictive accuracy under kernel ridge regression was also consistent across pipelines. Using the connectivity kernel, the Schaefer atlas consistently yielded the highest prediction accuracy in every pipeline, whereas the gradient-based method consistently yielded the lowest. In the structural equation models, the two gMSHBM parcel size paths were the only ones that reached significance in all four pipelines (*β* = .16 to .22). Every other significant path appeared under some pipelines but not others.

However, we found that other metrics were sensitive to preprocessing choices. Temporal filtering uniformly inflated functional homogeneity across all methods. Most notably, the total number of significant regression paths (out of the 126 tested) varied substantially across pipelines. Pipeline A yielded only 6 significant paths. In contrast, pipelines B, C, and D yielded 27, 63, and 47 significant paths, respectively. These additional paths were concentrated in the features derived from the time series rather than in those derived from the geometry of the maps. None of the six paths in pipeline A involved global efficiency or connectivity similarity, whereas these two features accounted for 17 of the 27 paths in pipeline B, 49 of the 63 in pipeline C, and 37 of the 47 in pipeline D. Temporal filtering and global signal regression act directly on the time series from which both features are computed, while the parcel size features remained nearly unchanged across pipelines. These results indicate that brain-behavior associations based on time-series-derived features were particularly sensitive to preprocessing choices, whereas associations involving parcel-size features were more stable.

## 4. Discussion

We evaluated how different algorithmic strategies for incorporating group-level, spatial, and individual information shape personalized cortical maps and relate to their predictive utility. Overall, the results revealed several recurring trade-offs across the methods evaluated. Methods with fewer explicit group-derived or spatial constraints showed higher functional homogeneity in our analyses, but also tended to show lower cross-individual spatial correspondence and session-to-session spatial stability. We also found that different methods influence the detection of candidate ectopic variants of the personalized parcels. Brain-behavior association analyses revealed a methodological divide. Topographical variation, specifically parcel size, was associated with general intelligence primarily when grounded in the shared spatial framework of gMSHBM. Yet, when predicting intelligence directly from functional connectivity matrices, personalization did not improve predictive accuracy relative to the group atlas. The standard group atlas yielded the highest prediction accuracy among the methods evaluated. Finally, by testing across distinct preprocessing pipelines, we found that analytical pipeline configuration particularly affected brain-behavior associations involving time-series-derived features, whereas relative between-method patterns in spatial characteristics were more stable. Because the methods differ in multiple algorithmic properties, these between-method patterns should not be interpreted as effects of a single underlying level of individualization or group constraint.

### 4.1. Algorithmic Constraints Shape the Intrinsic Profile of Personalized Maps

Four of the six evaluated approaches (Schaefer, gMSHBM, AGP, and SLIC-F) yielded similar functional homogeneity (.305 to .318), while all six produced comparable test-retest reliability (.842 to .867). For the four methods clustered in the lower homogeneity range, this convergence in homogeneity may reflect different forms of spatial or group-derived regularization that limit how freely parcel boundaries can adapt to individual functional similarity. The gradient-based method and SLIC-C achieved higher functional homogeneity (.379 and .373, respectively) both rely more directly on individual functional information and lack an explicit group-derived parcel reference during boundary estimation, providing greater flexibility to identify boundaries based on within-individual functional similarity. Meanwhile, the convergence in test-retest reliability across all methods might reflect the underlying data quality rather than algorithmic constraints. The reliability of resting-state connectivity depends heavily on scan length and signal-to-noise ratio (Birn et al., 2013; Vale et al., 2026). The HCP-YA protocol provides high-quality data acquired with a short repetition time (0.72 s), with 2,400 time points per session, supporting relatively stable connectivity estimates within each session. This finding aligns with previous work demonstrating the stability of HCP-YA single-session connectivity profiles (Finn et al., 2015), and the increased reliability obtained with longer scans (Noble et al., 2017; Vale et al., 2026). Thus, the high connectivity reliability observed across methods may partly reflect the favorable data quality and scan duration of the HCP-YA dataset, rather than indicating that parcellation constraints have little influence on reliability.

The inter-individual Dice coefficient also differed substantially across methods, consistent with differences in their algorithmic design and use of shared spatial information. Since it measures how much parcellations overlap across individuals, a lower value indicates lower cross-individual spatial correspondence. The Schaefer atlas scores a perfect value given that there is no variability across individuals. Among the personalized methods, approaches enforcing spatial or probabilistic constraints (SLIC-F, .687; gMSHBM, .554) produced higher overlap across individuals than methods retaining only seed locations (AGP, .443) or ignoring group priors entirely (gradient-based, .269; SLIC-C, .157). This ordering is broadly consistent with whether and how each algorithm incorporates a shared spatial reference, although the methods also differ in several other computational properties. A highly personalized map might reflect a genuine neurobiological profile, or it might also reflect sensitivity to measurement noise or transient functional states. Testing whether similar spatial organization re-emerges from a separate scan of the same individual provides a way to further evaluate this ambiguity. SLIC-F and AGP reached session-to-session Dice coefficients of .761 and .592, respectively, whereas the gradient-based method reached only .318 and SLIC-C .255. The two unconstrained methods (SLIC-C and the gradient-based method) showed lower recovery of their parcellations across sessions. This low spatial stability is consistent with greater sensitivity to transient noise, although it could also reflect sensitivity to genuine session-to-session changes in functional organization. Constrained methods showed higher cross-session spatial stability in our analyses. Distinguishing stable individual topography from transient variability therefore remains an open challenge. Resolving this ambiguity would benefit from densely sampled designs, where extensive repeated measurements can help disentangle stable spatial variation from transient state fluctuations or noise before extracting biological conclusions (Chang et al., 2026; Gordon et al., 2017; Laumann et al., 2015). At the same time, without shared spatial constraints, the parcels of unconstrained methods are difficult to match across individuals. How to establish such correspondence remains an open question in personalized fMRI, with potential solutions including task fMRI (Gratton & Braga, 2026) or multimodal data (Kristanto, 2026). Unconstrained methods therefore require additional validation across repeated measurements and complementary data to determine whether their individualized maps reflect stable topography rather than transient variation or noise.

Next, spatial contiguity provides information regarding the detection of candidate topographic variants in personalized parcels. Dworetsky et al. (2024) described these two forms of variation at the network level, but comparable variation also occurs at the areal level. For example, Yang et al. (2025) found an atypical topology of area 55b in 19.1% of left and 18.4% of right hemispheres among 1,071 HCP-YA participants, and architectonic data point in the same direction, as area hOc5 can appear away from its group-average location (Fischl et al., 2008). Our results show that the opportunity to detect such spatially disconnected components depends strongly on the parcellation method. The parcels derived from the Schaefer atlas were fully contiguous. AGP, SLIC-F, and the gradient-based method (Gordon et al., 2016) produced highly contiguous parcels because their algorithms enforce spatially adjacent assignments. In contrast, gMSHBM and SLIC-C produced highly fragmented parcels. gMSHBM explicitly allows non-contiguous parcels within its spatial localization constraints (Kong et al., 2021), while SLIC-C lacks spatial constraints entirely, resulting in substantially greater fragmentation.

Classifying a fragment as a candidate ectopic variant depended heavily on the chosen mathematical distance threshold. Under gMSHBM, 68.2% of fragments exceeded 3.5 mm, but only 0.7% exceeded 30 mm. The literature lacks a consensus threshold for this classification. Dworetsky et al. (2024) applied a 3.5 mm cutoff at the network level; importantly, their primary definition measured variant distance relative to the corresponding territory in a group-average network map, whereas we measured each secondary fragment relative to the main body of the corresponding personalized parcel. Their subsequent analyses showed that most group-defined ectopic variants also remained spatially isolated within individual-specific network maps, but the two operationalizations are not directly equivalent. Therefore, the 3.5 mm value in our analysis should be interpreted as a reference threshold rather than a directly transferable criterion for biological ectopy. In addition, because our parcels were defined at the areal rather than network level, the appropriate distance criterion may differ.

Validating these variants requires tasks that isolate the function of individual areas, which remains a central challenge in cognitive neuroscience (Kristanto et al., 2022; Poldrack & Yarkoni, 2016). Another challenge is that appropriate cutoffs may vary based on data quality, spatial smoothing, and registration accuracy (Alakörkkö et al., 2017). A threshold calibrated on high-quality HCP data may not transfer directly to clinical populations with higher motion or shorter scans (Kristanto et al., 2026; Pardoe et al., 2016). Spatial fragmentation alone does not establish biological validity; candidate variants should therefore be interpreted cautiously and, where possible, validated using independent sessions, behavioral measures, task activation, or complementary modalities before being considered evidence of neurobiological reorganization.

Finally, preprocessing choices altered the absolute values of several intrinsic metrics, while many relative between-method patterns were more stable. Implementing personalized parcellation therefore involves at least two distinct analytical decisions. The choice of algorithm influences the topology of the map through its specific objective function, initialization, and group or spatial constraints. Preprocessing steps, coupled with data quality, can further alter the estimated levels of homogeneity, reliability, contiguity, and downstream brain-behavior associations.

### 4.2. Personalized Topography and Connectivity Show Different Predictive Patterns

We found that topographical variation was associated with *g*, most consistently under gMSHBM. With gMSHBM, the intra-individual coefficient of variation for parcel size, which quantifies how unevenly functional boundaries expand or contract relative to the rest of the cortex, was positively associated with *g* (*β* = .22). No comparable association survived correction for any other method or feature. Unconstrained methods, such as SLIC-C and the gradient-based method, generated substantially greater variance in parcel sizes (coefficients of 1.027 and 1.890, respectively) consistent with their weaker spatial constraints. In contrast, the spatial localization prior in gMSHBM restricts boundary shifts to within 30 mm of the group average. Within this constrained framework, a high coefficient of variation indicates greater variation in the relative sizes of parcels while maintaining correspondence to a shared spatial reference. These findings align with a growing body of literature linking the spatial extent of functional territories to behavior. Keller et al. (2023) demonstrated a positive correlation between the total cortical representation of fronto-parietal networks and general cognition. Moreover, Anderson et al. (2021) showed that individualized network topography, quantified by the coefficient of variation of network size, is heritable and exhibits the greatest variability in the heteromodal cortex. Our findings, supported by previous studies, are consistent with the possibility that individual variation in the spatial extent of functional territories carries behaviorally relevant information.

When predicting *g* from functional connectivity matrices, however, the group atlas yielded higher predictive accuracy than the personalized methods evaluated here. Using the connectivity kernel, the static Schaefer group atlas predicted *g* better than every personalized method. The predictive performance of the personalized methods broadly followed their inter-individual Dice coefficients, with two exceptions: SLIC-F had the highest Dice among them but ranked fourth in accuracy, and the gradient-based method had a higher Dice than SLIC-C but the lowest accuracy of all. This raises the question of why functional connectivity derived from a group atlas with lower functional homogeneity showed higher predictive accuracy than connectivity derived from individualized maps.

One plausible explanation lies in the edge-wise correspondence required by the kernel ridge regression model. Kernel ridge regression predicts the g score of each individual by calculating how similar their full connectivity matrix is to those of other participants. Specifically, the algorithm evaluates this similarity edge by edge. Under a fixed group atlas, a specific edge refers to the same atlas-defined pair of parcels across participants. Under personalized methods, individual boundary shifts reduce exact spatial correspondence between nominally matched parcels. This mismatch may reduce the comparability of corresponding edges across individuals and thereby weaken the similarity structure used by the predictive model. The broad agreement between prediction accuracy and the Dice ranking is consistent with this interpretation. It is important to note that cross-individual correspondence among the personalized parcellations was established by one-to-one Hungarian alignment to the Schaefer atlas. Thus, parcel labels were assigned to maximize spatial overlap with this common reference, rather than being independently homologized across individuals. This procedure enabled parcel-wise and edge-wise comparisons, but it may also favor methods whose spatial organization remains closer to the Schaefer reference.

This interpretation is also consistent with the strong negative relationship between parcel-size variability and global efficiency across methods within individuals. Methods producing highly unequal parcel sizes may alter the representation of functional connectivity by averaging signals over parcels that differ substantially in spatial extent and functional composition. Such discretization effects could influence both graph-level measures and edge-wise predictive models. However, because parcel-size variability co-varies with other algorithmic properties, our analyses cannot determine whether parcel-size inequality itself causes the reduction in connectivity-based performance.

These results suggest different methodological trade-offs associated with distinct parcellation strategies in brain-behavior modeling. In our analyses, individualized spatial features were behaviorally informative under gMSHBM, which retains correspondence to a shared spatial reference. In contrast, prediction from edge-wise functional connectivity may benefit from stronger cross-individual spatial correspondence, as observed for the group atlas in our KRR analyses. Thus, the relative utility of personalization may depend on whether the downstream analysis relies primarily on individual topography or on directly comparable connectivity edges across participants.

### 4.3. Toward an Integrative Framework for Personalized Parcellation

Our results highlight multiple methodological trade-offs among personalized parcellation strategies. Algorithms differ not only in whether they incorporate group-derived information, but also in their objective functions, spatial constraints, initialization procedures, and treatment of disconnected regions. Group-derived or spatially restrictive information can help preserve cross-participant correspondence, but may also limit the expression of individual spatial variation. In contrast, approaches driven more directly by individual functional data can accommodate greater spatial variability, but in our analyses some of these methods showed lower cross-session stability or cross-individual correspondence. These patterns should not be interpreted as identifying an optimal level of individualization, because such a question would require manipulating the level of group or spatial constraints within the same algorithm while holding other design choices constant, such as hierarchical Bayesian models with adaptable group priors.

Evaluating these compromises remains a practical challenge. An integrative, interactive framework offers one way to address it (Kristanto, 2026). Such a platform would enable researchers to systematically explore methods across categories, tracking their empirical utilization, their performance across standardized evaluation criteria, and their documented computational requirements. This approach aligns with the METEOR application developed for preprocessing pipelines in graph-based fMRI (Kristanto et al., 2024) and similar frameworks for mobile EEG preprocessing (Jacobsen et al., 2025). These tools make the space of analytical options easier to navigate. Coupling this interactive framework with a unified computational toolbox (Burkhardt & Gießing, 2026), as implemented through the single codebase in this study, allows researchers to move from method selection to execution. The utility of this framework also extends beyond individual study design to support broader meta-scientific investigations. By systematically mapping the analytical space, researchers can identify prevalent techniques, recurring evaluation failures, and unresolved methodological gaps. For example, none of the methods evaluated here simultaneously combined the ability to represent spatially disconnected parcels with strict cross-individual correspondence, highlighting a methodological tension that warrants further development. Mapping these limitations systematically could help guide future algorithm development toward addressing documented methodological gaps rather than relying on individual intuition.

### 4.4. Limitations

This study presents four primary limitations. First, our sample consists exclusively of healthy young adults providing dense, high-quality resting-state data. Because functional reliability depends partly on scan length, the cross-method convergence in test-retest reliability observed here may not generalize directly to datasets characterized by shorter scan durations or greater head motion, especially in clinical populations. Second, we evaluated all methods at a fixed resolution of 200 parcels to guarantee mathematical comparability. Since functional homogeneity inherently depends on parcel count, our current data cannot establish whether these relative methodological rankings remain stable across other spatial resolutions. Third, to maximize reproducibility and strictly adhere to the reference algorithmic framework, we used the previously published set of gMSHBM parcellations. Because these specific boundaries rely on the fixed preprocessing choices of the original authors, they did not vary across our four experimental pipelines. Accordingly, our preprocessing comparison does not test how re-estimating gMSHBM under different preprocessing choices would affect its individualized boundaries. Instead, it evaluates how downstream measures derived from those fixed maps change when the associated time series are processed differently. Finally, another limitation concerns the operational definition of candidate ectopic variants. We measured fragment distance relative to the main body of each personalized parcel, whereas the 3.5 mm reference threshold from Dworetsky et al. (2024) was originally defined relative to a group-average network map. Although their later analyses showed that many such variants also remained isolated in individual-specific maps, these definitions are not equivalent. Our threshold analysis should therefore be interpreted as a methodological sensitivity analysis rather than as an estimate of the biological prevalence of ectopic variants.

## 5. Conclusion

Personalized parcellation methods impose distinct algorithmic constraints that shape the topology and downstream analytical properties of the resulting brain maps. No single algorithm performed optimally across all criteria evaluated here. Our findings therefore suggest that the consequences of method choice depend on the downstream analytical objective. First, for prediction from functional connectivity, the group atlas provided the highest accuracy among the methods evaluated here, suggesting that cross-individual spatial correspondence may be advantageous for edge-wise predictive models. Second, associations between spatial topography and behavior were most consistent for gMSHBM, where parcel size variability and similarity were associated with g while retaining correspondence to a shared spatial framework. This pattern suggests that individualized spatial features may be most interpretable when variation is measured relative to a common reference, although this requires further validation across methods and datasets. Third, the detection of candidate ectopic variants depends strongly on both algorithmic constraints and the chosen distance criterion. Methods that enforce contiguity cannot represent spatially disconnected components, whereas methods that permit disconnected components can capture such configurations. Determining whether these configurations reflect stable individualized topography rather than transient variability or noise requires independent validation., ideally using repeated measurements and complementary functional or multimodal evidence.

## Data Availability Statement

The HCP-YA data are available from the Human Connectome Project (https://www.humanconnectome.org/) under the HCP data use terms; access to the restricted data requires a separate application. The gMSHBM parcellations are available from https://github.com/ThomasYeoLab/CBIG. All analysis code is available at https://github.com/CoskuInceler/Individualized_Brain_Parcellation.

## CRediT authorship contribution statement

Yusuf Cosku Inceler: Conceptualization, Methodology, Software, Formal analysis, Visualization, Writing – original draft. Micha Burkhardt: Data curation, Resources, Writing – review & editing. Carsten Gießing: Conceptualization, Supervision, Funding acquisition, Writing – review & editing. Daniel Kristanto: Conceptualization, Methodology, Project administration, Supervision, Funding acquisition, Writing – original draft, Writing – review & editing.

## Acknowledgment

This work was supported by the Young Researchers’ Fellowship from the University of Oldenburg awarded to DK and by funding from the German Research Foundation (Deutsche Forschungsgemeinschaft; DFG) to Carsten Gießing (Project No. 464552782; GI 682/5-2) as part of the DFG Priority Program “META-REP: A Metascientific Program to Analyze and Optimize Replicability in the Behavioral, Social, and Cognitive Sciences” (SPP 2317). This work was also partly supported by the DFG under Germany’s Excellence Strategy – EXC 2177/1 – Project ID 390895286. Data were provided by the Human Connectome Project, WU-Minn Consortium (Principal Investigators: David Van Essen and Kamil Ugurbil; 1U54MH091657), funded by the 16 NIH Institutes and Centers that support the NIH Blueprint for Neuroscience Research, and by the McDonnell Center for Systems Neuroscience at Washington University.

