## Supplemental Materials for "From Intrinsic Topography to Cognitive Prediction: Benchmarking Approaches to Personalized Cortical Parcellation"

### Supplementary Materials

#### Supplementary Material A. Results Under the Remaining Preprocessing Pipelines

The main text presents the results of pipeline A, which adds no cleaning steps beyond those already performed by the HCP pipelines. Tables A1 and A2 provide the corresponding results for pipelines B, C, and D.

Two patterns are noteworthy. Temporal filtering increased functional homogeneity by .12 to .14 for each method and reduced the differences between methods; the range across methods narrowed from .074 in pipeline A to .067 in pipeline B, while all values increased. Filtering also reduced contiguity for the two SLIC variants, from 31.2 to 8.7% for SLIC-C and from 99.6 to 94.5% for SLIC-F, whereas the other methods were unaffected. Global signal regression slightly reduced homogeneity and did not alter the ranking of the methods.

The inter-individual Dice coefficient remained nearly unchanged across pipelines for all methods except SLIC-F, where it decreased from .687 in pipeline A to .553 in pipeline B. SLIC-F is the only method whose compactness parameter was recalibrated per pipeline, and filtering altered the distribution from which the calibration was derived.

**Table A1.** *Validation metrics under the remaining three preprocessing pipelines.* Mean (*SD*) across 947 participants. Conventions as in Table E2.

| Pipeline | Method | Homogeneity | Test-retest | Contiguity (%) | Dice |
|---|---|---|---|---|---|
| B (filtered) | Schaefer | .439 (.042) | .859 (.052) | 100.00 (0.00) | 1.000 |
| | gMSHBM | .457 (.044) | .862 (.052) | 61.51 (11.19) | .554 (.014) |
| | AGP | .459 (.043) | .855 (.052) | 100.00 (0.00) | .440 (.008) |
| | SLIC-F | .448 (.045) | .852 (.055) | 94.45 (2.43) | .553 (.010) |
| | SLIC-C | .494 (.042) | .797 (.067) | 8.65 (2.70) | .157 (.012) |
| | Gradient-based | .506 (.035) | .839 (.052) | 99.99 (0.07) | .270 (.014) |
| C (minimal + GSR) | Schaefer | .288 (.023) | .862 (.054) | 100.00 (0.00) | 1.000 |
| | gMSHBM | .298 (.026) | .868 (.054) | 61.51 (11.19) | .554 (.014) |
| | AGP | .297 (.024) | .862 (.054) | 100.00 (0.00) | .452 (.007) |

| Pipeline | Method | Homogeneity | Test-retest | Contiguity (%) | Dice |
|---|---|---|---|---|---|
| | SLIC-F | .283 (.024) | .856 (.056) | 99.69 (0.40) | .694 (.016) |
| | SLIC-C | .352 (.023) | .840 (.060) | 26.86 (5.12) | .157 (.010) |
| | Gradient-based | .366 (.022) | .858 (.054) | 99.99 (0.07) | .273 (.015) |
| D (filtered + GSR) | Schaefer | .411 (.033) | .847 (.050) | 100.00 (0.00) | 1.000 |
| | gMSHBM | .430 (.038) | .852 (.050) | 61.51 (11.19) | .554 (.014) |
| | AGP | .433 (.036) | .843 (.051) | 100.00 (0.00) | .447 (.008) |
| | SLIC-F | .419 (.037) | .838 (.054) | 95.02 (2.29) | .558 (.011) |
| | SLIC-C | .472 (.036) | .780 (.067) | 8.05 (2.62) | .153 (.011) |
| | Gradient-based | .490 (.032) | .839 (.048) | 99.99 (0.06) | .272 (.015) |

Table A2. *Prediction of general intelligence under the remaining three preprocessing pipelines. Conventions as in Table 3;* * FDR-corrected $p < .05$.

| Pipeline | Method | KRR $r$ (conn.) | KRR $r$ (size) | SEM $R^2$ | Unique $R^2$ (size) |
|---|---|---|---|---|---|
| B (filtered) | Schaefer | .382** | — | .0207 | — |
| | gMSHBM | .350** | .218** | .0613 | .0457 |
| | AGP | .316** | .058 | .0189 | .0096 |
| | SLIC-F | .297** | .076* | .0099 | .0004 |
| | SLIC-C | .321** | .052 | .0428 | .0254 |
| | Gradient-based | .162** | −.048 | .0079 | .0002 |
| C (minimal + GSR) | Schaefer | .391** | — | .0472 | — |
| | gMSHBM | .377** | .218** | .0798 | .0217 |
| | AGP | .365** | .042 | .0395 | .0008 |
| | SLIC-F | .340** | .007 | .0613 | .0114 |
| | SLIC-C | .272** | .104** | .0145 | .0004 |

| Pipeline | Method | KRR *r* (conn.) | KRR *r* (size) | SEM $R^2$ | Unique $R^2$ (size) |
|---|---|---|---|---|---|
| | Gradient-based | .164** | −.034 | .0349 | .0024 |
| D (filtered + GSR) | Schaefer | .403** | — | .0316 | — |
| | gMSHBM | .350** | .218** | .0637 | .0331 |
| | AGP | .332** | .101** | .0251 | .0068 |
| | SLIC-F | .300** | .101** | .0221 | .0026 |
| | SLIC-C | .327** | .056 | .0278 | .0142 |
| | Gradient-based | .097** | −.036 | .0207 | .0016 |

The gMSHBM parcel size kernel yielded exactly $r = .218$ in all four pipelines. This constancy does not indicate that gMSHBM is robust to preprocessing. The gMSHBM parcellations were precomputed by the original authors and are identical in every pipeline, so features that depend only on parcel sizes cannot change. The connectivity kernel for the same method, which depends on the preprocessed time series, did vary ($r = .350$ to .377).

### Supplementary Material B. The Gradient-Based Method at Its Native Resolution

In the main text, all methods were compared at 200 parcels. Reducing the gradient-based method to this number alters it, and we report here its behavior at the resolution implied by its boundary map.

Across 947 participants, pipeline A produced 603.3 (*SD* 21.6) parcels, against the 620.8 (*SD* 39.4) reported by the original authors on the Midnight Scan Club dataset (Gordon et al., 2017). This agreement supports the fidelity of the reimplementation. Parcel counts were slightly lower under the other pipelines (590.9 in pipeline B, 591.8 in pipeline C, and 577.7 in pipeline D).

At its native resolution the method achieved a functional homogeneity of .458 (*SD* .028), against .379 at 200 parcels, which is the expected consequence of smaller parcels. Test-retest reliability was .840 (*SD* .066), slightly lower than the .852 obtained at 200. Contiguity remained at 100%, and the inter-individual Dice coefficient reached .470, considerably higher than the .269 obtained after merging, indicating that the merging step reduces the correspondence of parcels across participants.

The method assigned 70.8% (*SD* 0.4) of cortical vertices, leaving the remainder on strong boundaries as transition zones. This is a design decision of the original method rather than a failure of convergence, and it is one reason its homogeneity is high: the vertices it declines to assign are those that would otherwise reduce it.

#### Supplementary Material C. Verification of the Reimplementations

The three algorithms that we ran ourselves (AGP, SLIC, and the gradient-based method) were reimplemented within a single `Python` codebase. Since a reimplementation might differ from the original in ways not apparent in the output, each was validated by comparing it to a direct implementation of the published formulation.

##### C.1. Computational Reformulations

Two steps were reformulated to make the entire sample processable. In both cases, the reformulation is algebraically equivalent to the original.

In the region growing step of AGP, the average correlation between a candidate vertex and the current members of a parcel is obtained from the running sum of the members' z-scored time series, rather than by constructing the entire member-by-candidate correlation matrix at each step. For a parcel P with members *m* and a candidate vertex v, the average correlation is given by

$$a(v, P) = 1/(|P| \cdot T) \cdot z_v \cdot \sum_{(m \in P)} z_m$$

where *z* represents a z-scored time series and *T* the number of time points. The cumulative sum is updated incrementally as vertices are added to the parcel.

In SLIC, the squared distance between a vertex v and a cluster center c is expanded as follows:

$$\|v - c\|^2 = \|v\|^2 - 2v \cdot c + \|c\|^2$$

This reduces the assignment step to a single matrix multiplication performed simultaneously over all vertices and centers.

Each reformulation was validated by comparison with a direct implementation on a test participant. Both reproduced the original labels exactly.

##### C.2. Numerical Precision

There is one numerical detail worth noting. Calculating the SLIC distances in single precision resulted in parcellations that differed from the direct formulation at approximately 1% of vertices. The expansion above subtracts quantities of similar magnitude from one another, and the resulting loss of precision occasionally reverses an assignment, an effect that accumulates over iterations. In double precision the two implementations coincided exactly. Distances were therefore accumulated in double precision throughout.

##### C.3. Calibration of the SLIC Compactness Parameter

The compactness parameter *m* balances the functional and spatial terms in the SLIC distance. Two adjustments were made relative to the formulation used in previous work.

First, the functional distance was divided by √T, which makes it equal to √(2(1 − r)) and thus independent of the number of time points. Without this, a parcellation estimated from one session and one estimated from both sessions lie on different scales, and the same value of *m* carries a different meaning in each.

Second, the reference distribution was changed. When sampling random vertex pairs, the distances cluster around √2, because nearly any two vertices are uncorrelated with one another: across $10^6$ pairs, the median was 1.411 with a standard deviation of .028. Setting *m* to that median leaves the ratio $d_{func}/m$ at approximately 1 for every pair, so the functional term contributes almost no variation and the spatial term becomes the determining factor.

Restricting the sample to neighboring vertices, which is what the algorithm compares when determining where a vertex belongs, yielded a median of .822 and a standard deviation of .188, seven times the dispersion. This mirrors the original formulation, in which the color distance that determines the scale is the distance between adjacent pixels.

A sensitivity analysis confirmed that *m* controls the balance as intended. Lowering *m* below the calibrated value degraded SLIC-F toward SLIC-C: contiguity dropped to 94% at $m = 0.5$, to 63% at $m = 0.2$, and to 23% at $m = 0.05$, while homogeneity increased in parallel. The calibrated value lies on the flat portion of this curve, which is why recalibration shifted SLIC-F toward individualization without altering its position in the comparison.

### Supplementary Material D. Bifactor Measurement Model

The measurement model was fitted to the nine cognitive tasks alone, with no brain data present, and the unstandardized loadings were then fixed for all subsequent models. The fit was acceptable: CFI = .970, TLI = .950, RMSEA = .051 [.038, .065], SRMR = .032, with standard errors clustered at the family level across 418 families and 940 participants.

**Table D1.** *Unstandardized loadings from the bifactor measurement model.*

| Factor | Indicator | Estimate | SE | *z* | *p* |
|---|---|---|---|---|---|
| *g* | Dimensional Change Card Sort | .313 | .040 | 7.83 | < .001 |
| *g* | Flanker | .231 | .043 | 5.41 | < .001 |
| *g* | Pattern Completion Processing Speed | .227 | .041 | 5.48 | < .001 |
| *g* | Picture Vocabulary | .605 | .044 | 13.68 | < .001 |
| *g* | Oral Reading Recognition | .645 | .042 | 15.40 | < .001 |
| *g* | Penn Progressive Matrices | .685 | .035 | 19.41 | < .001 |

| Factor | Indicator | Estimate | SE | z | p |
|---|---|---|---|---|---|
| *g* | Line Orientation | .588 | .042 | 14.04 | < .001 |
| *g* | Penn Word Memory | .307 | .042 | 7.34 | < .001 |
| *g* | Picture Sequence Memory | .333 | .045 | 7.37 | < .001 |
| speed | Dimensional Change Card Sort | .689 | .041 | 16.69 | < .001 |
| speed | Flanker | .640 | .044 | 14.71 | < .001 |
| speed | Pattern Completion Processing Speed | .473 | .043 | 10.99 | < .001 |
| crystallized | Picture Vocabulary | .528 | .031 | 16.93 | < .001 |
| crystallized | Oral Reading Recognition | .528 | .031 | 16.93 | < .001 |
| memory | Penn Word Memory | .327 | .054 | 6.01 | < .001 |
| memory | Picture Sequence Memory | .327 | .054 | 6.01 | < .001 |

The loadings within the crystallized and memory factors were constrained to be equal because each had only two indicators. All nine tasks exhibited a significant loading on *g*; the highest loadings were observed in the reasoning and crystallized tasks, while the speed tasks, whose variance was largely explained by their own group factor, were among the lowest. This is the pattern expected from a bifactor model of this test battery.

#### Supplementary Material E. Pairwise Comparisons Between Methods

The differences between each pair of methods were tested using paired *t*-tests, with the Holm correction applied across the 15 pairs within each measure, and reported as Cohen's $d_{av}$. At this sample size every comparison reached statistical significance, so the effect sizes are the quantity of interest.

**Table E1.** *Pairwise differences in functional homogeneity.* A positive *d* indicates that the first method scores higher. All comparisons $p < .001$ after Holm correction.

| Comparison | $d_{av}$ |
|---|---|
| Schaefer vs gMSHBM | −0.30 |
| Schaefer vs AGP | −0.23 |
| Schaefer vs SLIC-F | 0.13 |

| Comparison | $d_{av}$ |
|---|---|
| Schaefer vs SLIC-C | −2.20 |
| Schaefer vs Gradient-based | −2.47 |
| gMSHBM vs AGP | 0.07 |
| gMSHBM vs SLIC-F | 0.42 |
| gMSHBM vs SLIC-C | −1.82 |
| gMSHBM vs Gradient-based | −2.07 |
| AGP vs SLIC-F | 0.35 |
| AGP vs SLIC-C | −1.94 |
| AGP vs Gradient-based | −2.20 |
| SLIC-F vs SLIC-C | −2.27 |
| SLIC-F vs Gradient-based | −2.53 |
| SLIC-C vs Gradient-based | −0.22 |

The comparisons fell into two groups. The differences among the four constrained methods were small, ranging from 0.07 to 0.42, despite all reaching statistical significance. The differences between these four methods and the two unconstrained methods were an order of magnitude larger, ranging from 1.82 to 2.53. The difference between the two unconstrained methods was again small (0.22). Functional homogeneity therefore distinguishes constrained from unconstrained methods, but does little to separate the methods within either group.

**Table E2.** *Validation metrics by method under pipeline A.* Mean (*SD*) across 947 participants. Homogeneity is computed on the full time series. Contiguity is the percentage of parcels forming a single connected component. Inter-individual Dice is the mean overlap of a participant's parcellation with all 946 others, measured through vertex co-assignment rather than parcel identity; the Schaefer value is 1.000 by construction. All methods are compared at 200 parcels.

| Method | Parcels | Homogeneity | Test-retest | Contiguity (%) | Inter-individual Dice |
|---|---|---|---|---|---|
| Schaefer | 200 | .309 (.030) | .863 (.060) | 100.00 (0.00) | 1.000 (.000) |
| gMSHBM | 200 | .318 (.032) | .867 (.060) | 61.51 (11.19) | .554 (.014) |

| Method | Parcels | Homogeneity | Test-retest | Contiguity (%) | Inter-individual Dice |
|---|---|---|---|---|---|
| AGP | 200 | .316 (.031) | .862 (.061) | 100.00 (0.00) | .443 (.008) |
| SLIC-F | 200 | .305 (.032) | .859 (.062) | 99.64 (0.46) | .687 (.016) |
| SLIC-C | 200 | .373 (.028) | .842 (.063) | 31.22 (5.26) | .157 (.010) |
| Gradient-based | 200 | .379 (.027) | .852 (.061) | 99.99 (0.08) | .269 (.015) |

### Supplementary Material F. Regression Paths Under the Two Binary Graph Thresholds

Table 2 in the main text reports the structural equation model estimates for the weighted graph. The table below reports the same models for the two binary graph thresholds, the graph preserving the strongest 10% of positive edges and the graph preserving edges with $r > .5$. The estimates are stable across thresholds: the coefficient of variation of gMSHBM parcel sizes and gMSHBM parcel size similarity remain the only paths that survive correction, and no other path reaches significance under either threshold.

| Threshold | Method | Predictor set | FC similarity | Global efficiency | Parcel size CV | Parcel size similarity |
|---|---|---|---|---|---|---|
| Strongest 10% | Schaefer | none | .100 | −.016 | — | — |
| | gMSHBM | CV | .067 | .009 | .220*** | — |
| | gMSHBM | similarity | .023 | .021 | — | .171*** |
| | gMSHBM | none | .079 | .009 | — | — |
| | AGP | CV | .043 | .029 | .066 | — |
| | AGP | similarity | .031 | .019 | — | .058 |
| | AGP | none | .050 | .022 | — | — |
| | SLIC-F | CV | .068 | .007 | −.065 | — |
| | SLIC-F | similarity | .075 | .008 | — | −.028 |
| | SLIC-F | none | .068 | .008 | — | — |
| | SLIC-C | CV | .027 | −.055 | −.021 | — |
| | SLIC-C | similarity | .036 | −.048 | — | −.008 |

| | | | | | | |
|---|---|---|---|---|---|---|
| | SLIC-C | none | .035 | −.048 | — | — |
| | Gradient-based | CV | .030 | .016 | −.007 | — |
| | Gradient-based | similarity | .031 | .015 | — | .016 |
| | Gradient-based | none | .033 | .015 | — | — |
| $r > .5$ | Schaefer | none | .096 | .056 | — | — |
| | gMSHBM | CV | .068 | .040 | .214*** | — |
| | gMSHBM | similarity | .018 | .117 | — | .200*** |
| | gMSHBM | none | .081 | .071 | — | — |
| | AGP | CV | .047 | −.001 | .064 | — |
| | AGP | similarity | .023 | .044 | — | .058 |
| | AGP | none | .042 | .046 | — | — |
| | SLIC-F | CV | .063 | .034 | −.067 | — |
| | SLIC-F | similarity | .070 | .022 | — | −.019 |
| | SLIC-F | none | .064 | .029 | — | — |
| | SLIC-C | CV | .043 | −.026 | −.010 | — |
| | SLIC-C | similarity | .045 | −.020 | — | −.006 |
| | SLIC-C | none | .044 | −.020 | — | — |
| | Gradient-based | CV | .026 | .108 | −.006 | — |
| | Gradient-based | similarity | .028 | .107 | — | .013 |
| | Gradient-based | none | .029 | .108 | — | — |

**Table F1.** Standardized regression paths from brain features to *g* under the two binary graph thresholds. Conventions as in Table 2. Asterisks denote significance after false discovery rate correction across all 126 regression paths (***$p < .001$). Dashes mark predictors not included in that set or unavailable for that method.
